# The Ketogenic Diet Metabolite β-Hydroxybutyrate Promotes Mitochondrial Elongation via Deacetylation in HeLa Cells and Improves Autism-like Behavior in Zebrafish

**DOI:** 10.1101/2022.10.03.510695

**Authors:** Golam M. Uddin, Rachel Lacroix, Mashiat Zaman, Iman Al Khatib, Amit Jaiswal, Jong M. Rho, Deborah M. Kurrasch, Timothy E. Shutt

## Abstract

The ketogenic diet (KD) is clinically beneficial and has therapeutic potential across a growing list of neurological disorders, including autism spectrum disorder (ASD). However, the underlying mechanisms mediating the benefits of the KD, which can also have undesirable side effects, remain undefined. To this end, improvements in mitochondrial morphology and function correlate with improved ASD behaviours in response to the KD, though how the KD influences mitochondrial morphology, and whether this is sufficient to improve behaviour remains unknown. Here, we investigate how beta-hydroxybutyrate (ΒHB), a key metabolite produced by the KD regulates mitochondrial morphology in HeLa cells, and whether this pathway could be exploited to alter phenotypes in a zebrafish model of ASD. We found that β-oxidation of ΒHB promotes mitochondrial elongation in HeLa cells by increasing NAD^+^ levels, which in turn activates SIRT deacetylases that act on key regulators of both mitochondrial fusion and fission. Our data suggest that increasing NAD^+^ levels with its precursor, nicotinamide mononucleotide (NMN), is sufficient to promote mitochondrial hyperfusion. Finally, both ΒHB and NMN impact neurodevelopment in the *shank3b*^+/−^ zebrafish model of ASD. Together, our findings elucidate a mechanism by which the ketogenic diet promotes mitochondrial elongation. Moreover, manipulation of this pathway may provide a novel avenue for the treatment of neurological disorders such as ASD that also may obviate potential complications of the KD in clinical practice.

## Introduction

The high-fat low-carbohydrate ketogenic diet (KD), long appreciated for its clinical efficacy for epilepsy (Neal et al., 2008), is also recognized to have potential beneficial effects in a growing list of diseases. The KD appears beneficial in neurodevelopmental disorders such as autism spectrum disorder (ASD) (Gogou & Kolios, 2018; Napoli et al., 2014), as well as in neurodegenerative diseases such as amyotrophic lateral sclerosis (Zhao et al., 2006), Parkinson disease and Alzheimer disease (Kashiwaya et al., 2000). For example, in the 3xTgAD mouse model of Alzheimer disease, the KD or ketone esters ameliorated AD pathogenesis, improved cognitive function and reduced anxiety. Moreover, due to its ability to manipulate mitochondrial energy metabolism, the KD may also be effective against cardiovascular diseases (Lopaschuk et al., 2020), or as a cancer treatment (Weber et al., 2020). Despite the well-documented benefits of the KD, concerns about limitations and complications of the diet are worth noting. For example, compliance with the KD can be challenging, especially in children and adolescents (Lin et al., 2017), with gastrointestinal side effects (39%) such as nausea/vomiting (28%), diarrhea (33%), constipation (2%), as well as hypertriglyceridemia (27%) being reported in patients (Kang et al., 2004). Understanding how the KD works will facilitate the development of alternative approaches that could avoid these complications.

Mitochondria are thought to be a central target mediating the effects of the KD. A key aspect regulating mitochondrial function is their dynamic structure, as they capable of fusing, dividing, and moving about the cell (Shutt & McBride, 2013; Wai & Langer, 2016). These processes influence mitochondrial function, with fragmentation of the mitochondrial network often associated with dysfunction (Baker et al., 2019; Glancy et al., 2020). Notably, recent work has shown that the KD can improve mitochondrial structure and function in the BTBR mouse model of ASD (Ahn et al., 2020), which is consistent with behavioral benefits in mouse models of ASD (Ahn et al., 2014; Ruskin et al., 2017; Ruskin et al., 2013), and clinical trials in children (El-Rashidy et al., 2017; Evangeliou et al., 2003; R. W. Y. Lee et al., 2018). However, the underlying mechanism(s) by which KD regulates mitochondrial structure remain unclear.

The ketone body β-hydroxybutyrate (ΒHB) is a key metabolic product of the KD that is thought to mediate many of the therapeutic effects of the KD (Newman & Verdin, 2017). There are multiple ways through which ΒHB is proposed to act, including inhibition of histone deacetylases that can elicit changes in gene expression (Shimazu et al., 2013), acting as a signaling molecule via G-protein-coupled receptors (GPCR) (Miyamoto et al., 2019), and as a metabolite consumed by mitochondria to generate ATP. With respect to this latter mechanism, when mitochondria catabolise ΒHB via β-oxidation instead of glucose, more NAD^+^ is available, with less being converted to NADH (Elamin et al., 2017; Newman & Verdin, 2017). Given that NAD^+^ is a necessary cofactor in several important enzymatic reactions, ΒHB-mediated changes in NAD^+^ can impact many cellular and mitochondrial functions. For example, NAD^+^ activates the sirtuin deacetylase enzymes (SIRTs) (Bonkowski & Sinclair, 2016; Covarrubias et al., 2021), which is a notable phenomenon since protein deacetylation can mediate mitochondrial elongation (Uddin et al., 2021). Indeed, an early report linking BHB to NAD+ metabolism found a high-fat diet was beneficial in a model of premature aging due elevated levels of ΒHB and activation SIRT1 (Scheibye-Knudsen et al., 2014) While NAD+ augmentation mirrored the benefits of the high fat diet, a link between BHB and increased NAD+ levels was not established at that time.

Mitochondrial dynamics, encompassing fusion, fission, mitophagy, and interactions with other organelles, are key aspects of maintaining mitochondrial function (Uddin, Abbas, & Shutt, 2021). Fusion and fission events allow mitochondria to adapt to the cell’s metabolic demands and impact mitochondrial morphology, cristae structure, energy production, apoptosis susceptibility, and mitochondrial quality control. Notably, NAD+ supplementation promotes mitophagy, which is crucial for mitochondrial maintenance (Fang et al., 2014; Gilmour et al., 2020). Meanwhile, NAD+ is a critical metabolite in the brain, where we also know mitochondrial function and mitochondrial dynamics are important, as NAD+ depletion is implicated in a growing list of neurodegenerative disorders (Fang et al., 2019). Understanding the intricate interplay between NAD+, protein acetylation, and mitochondrial dynamics is crucial for unraveling the mechanisms underlying mitochondrial dysfunction and exploring potential therapeutic interventions for related diseases.

Zebrafish are a valuable model for studying complex brain disorders such as ASD (Kalueff et al., 2014). Recently, genetic mutant models of verified autism-risk genes have been created and are shown effectively recapitulate autism-like behaviours (Durand et al., 2007; Liu et al., 2018). *SHANK3* encodes a master scaffolding protein that has critical roles in synaptogenesis and synaptic function (Monteiro & Feng, 2017), and is a validated ASD-risk gene in humans (SFARI, 2022; Yi et al. 2016). Moreover, *shank3b^+/−^* deficient zebrafish display core features of ASD-like behaviour, such as impaired locomotor activity, repetitive behaviours and social defects (S. Lee et al., 2018; Liu et al., 2018). Linking SHANK3 to mitochondria and mitochondrial function, mouse models show that the SHANK3 protein interacts with several mitochondrial proteins (Lee et al., 2017). Meanwhile, mice harboring a *Shank3* mutation associated with ASD had elevated S-nitrosylation of mitochondrial proteins, indicative of mitochondrial dysfunction (Kartawy et al., 2021).

While ΒHB treatment leads to mitochondrial elongation in cultured cells (Ahn et al., 2020; Santra et al., 2004), it is neither clear how ΒHB regulates mitochondrial morphology, nor whether ΒHB alone is sufficient to recapitulate the beneficial effects of the KD on ASD behaviours. Here, we investigated if and how mitochondrial morphology in HeLa cells is impacted by ΒHB-mediated changes in NAD^+^, and whether this pathway could potentially be exploited to improve behaviors in a zebrafish model of ASD.

## Results

### ΒHB mediated increase in NAD^+^ is necessary to promote mitochondrial elongation in HeLa cells

Building on previous work showing that BHB induces mitochondrial elongation in primary neuronal cultures (Ahn et al., 2020), we wanted to examine the mechanistic underpinnings of BHB-induced mitochondrial elongation in HeLa cells, a standard cell culture model that is often used to study mitochondrial dynamics as it is easy to work with and to visualize mitochondrial morphology. An initial dose response showed that a 24 hr treatment with BHB induces mitochondrial elongation in HeLa cells (S1). We chose to use the 5 mM dose as our treatment paradigm since this dose had the strongest effect on morphology and is a physiologically relevant dose for ketone levels found with the ketogenic diet. A more detailed analysis of mitochondrial morphology showed that 24 hr treatment with 5 mM ΒHB promoted mitochondrial elongation in HeLa cells (Figure 1A), as assessed by both quantitative analysis of mitochondrial branch length (Figure 1B), and qualitative analysis of overall mitochondrial morphology via visual scoring (Figure 1C). In conjunction with mitochondrial elongation, BHB treatment also increased basal oxygen consumption (S2). As ΒHB is reported to increase NAD^+^ via β-oxidation (Elamin et al., 2017), we measured levels of NAD^+^ in ΒHB-treated cells. As expected, treatment with ΒHB increased the levels of NAD^+^ (Figure 1D, S3B and S4D) and NAD^+^+ NADH levels (S3C and S4E). Notably, blocking β-oxidation using the CPT1a inhibitor etomoxir (Xu et al., 2003) prevented both ΒHB-mediated increases in NAD^+^ (Figure 1D) and mitochondrial elongation (Figure 1A-C, S4A-C), suggesting that β-oxidation is required for ΒHB to increase NAD^+^ levels, and that increased NAD^+^ is required for ΒHB-induced mitochondrial elongation.

**Figure 1:**
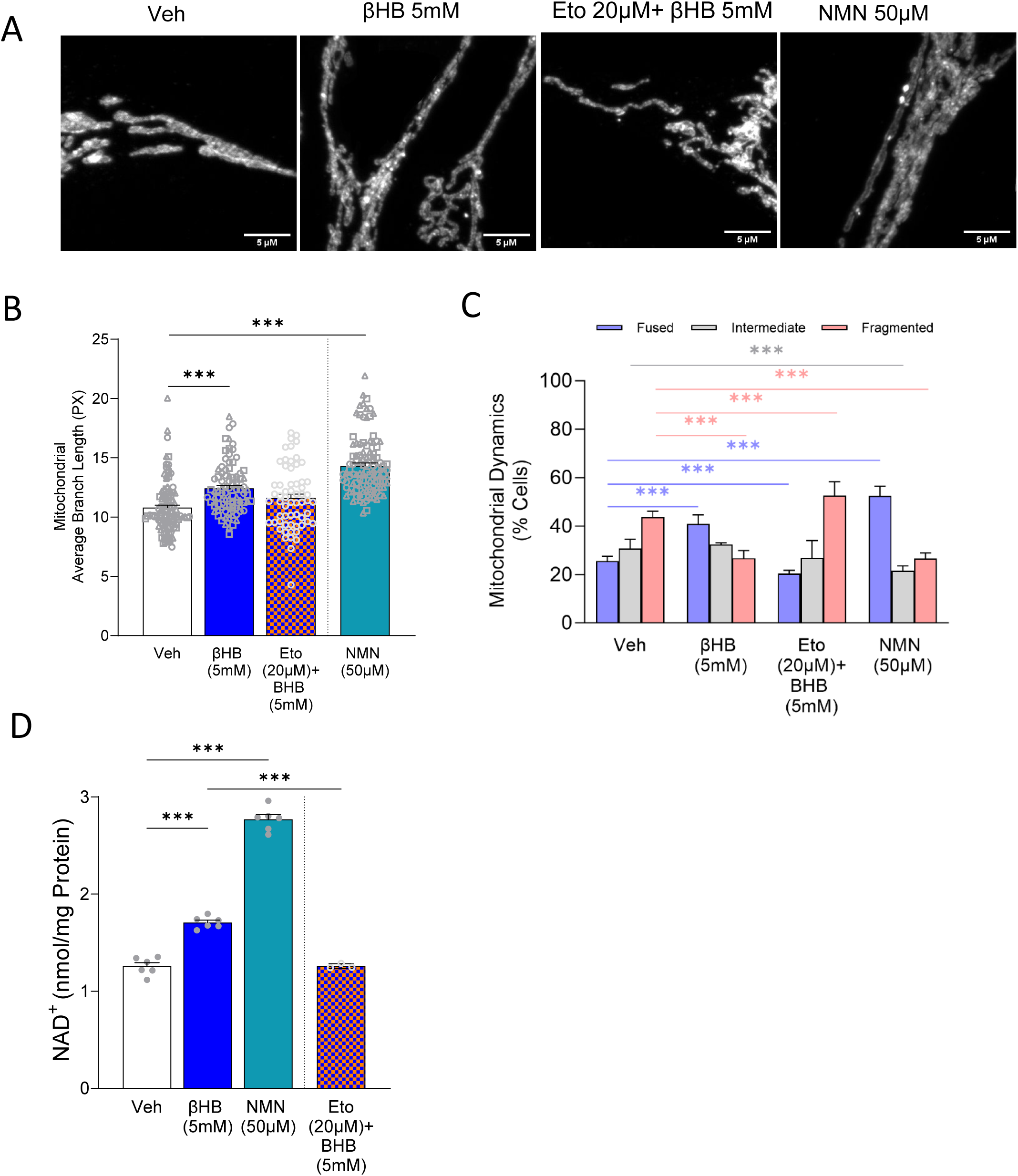
BHB and NMN promote mitochondrial elongation and increases NAD^+^ in Hela cell. **(A)** Representative confocal images of HeLa cells stained with antibodies against TOMM20 for mitochondrial network. HeLa cells were immuno-stained after 24h treatment with indicated concentrations of Veh or BHB or NMN or Eto+BHB. (B) Quantification of mean mitochondrial branch length were measured between Veh, BHB or NMN or Eto+BHB treated cells. This quantification was performed by Image-J, MiNA plugin from three independent replicates, using a standard region of interest (ROI) and measured randomly from each image. (C) Qualitative analysis of mitochondrial morphology was performed in Veh, BHB or NMN or Eto+BHB treated cells via an unbiased visual scoring method. Analysis was done using at least 150 cells per condition, and 3 independent replicates for each treatment condition. Statistical differences within the individual categories are colour-coded (blue = fused, gray = intermediate, and pink = fragmented) (D) NAD+ level was measured 24h post treatment of Veh or BHB or NMN; and Veh or Eto+BHB in HeLa cells (n=3-6/group). Data are shown as mean ± SEM. Data were analyzed using t-test. The significant differences are presented as *** p < 0.001.

### Increased NAD^+^ is sufficient to promote mitochondrial elongation

To investigate whether increased NAD^+^ alone could promote mitochondrial elongation, we treated cells with the NAD^+^ precursor molecule nicotinamide mononucleotide (NMN). Similar to the ΒHB treatment, we found that the NMN can increase both NAD^+^ levels (Figure 1D) and NAD^+^+ NADH levels (S3C), as well as promote mitochondrial elongation (Figure 1A-C).

### ΒHB and NMN promote mitochondrial elongation by increasing mitochondrial fusion and decreasing mitochondrial fission

Mitochondrial elongation can be caused by either increased fusion and/or decreased fission (Sabouny & Shutt, 2020). To investigate how ΒHB and NMN promote mitochondrial elongation, we examined key proteins mediating both mitochondrial fusion and fission (Figure 2A-D, S3D-G). We found that expression of the OPA1 fusion protein was increased significantly by treatment with ΒHB, but not NMN. Meanwhile, both ΒHB and NMN treatments reduced phosphorylation of DRP1 at position serine-616, which is a well-characterized post-translational modification that promotes mitochondrial fission (Sabouny & Shutt, 2020; Uddin et al., 2021). Notably, we did not see any significant changes due to ΒHB or NMN treatment in DRP1 phosphorylation at serine-637, which inhibits mitochondrial fission (Sabouny & Shutt, 2020; Uddin et al., 2021), nor in the total expression of other key mitochondrial fusion (MFN1, MFN2) or fission (MFF) proteins (S3D-G).

**Figure 2:**
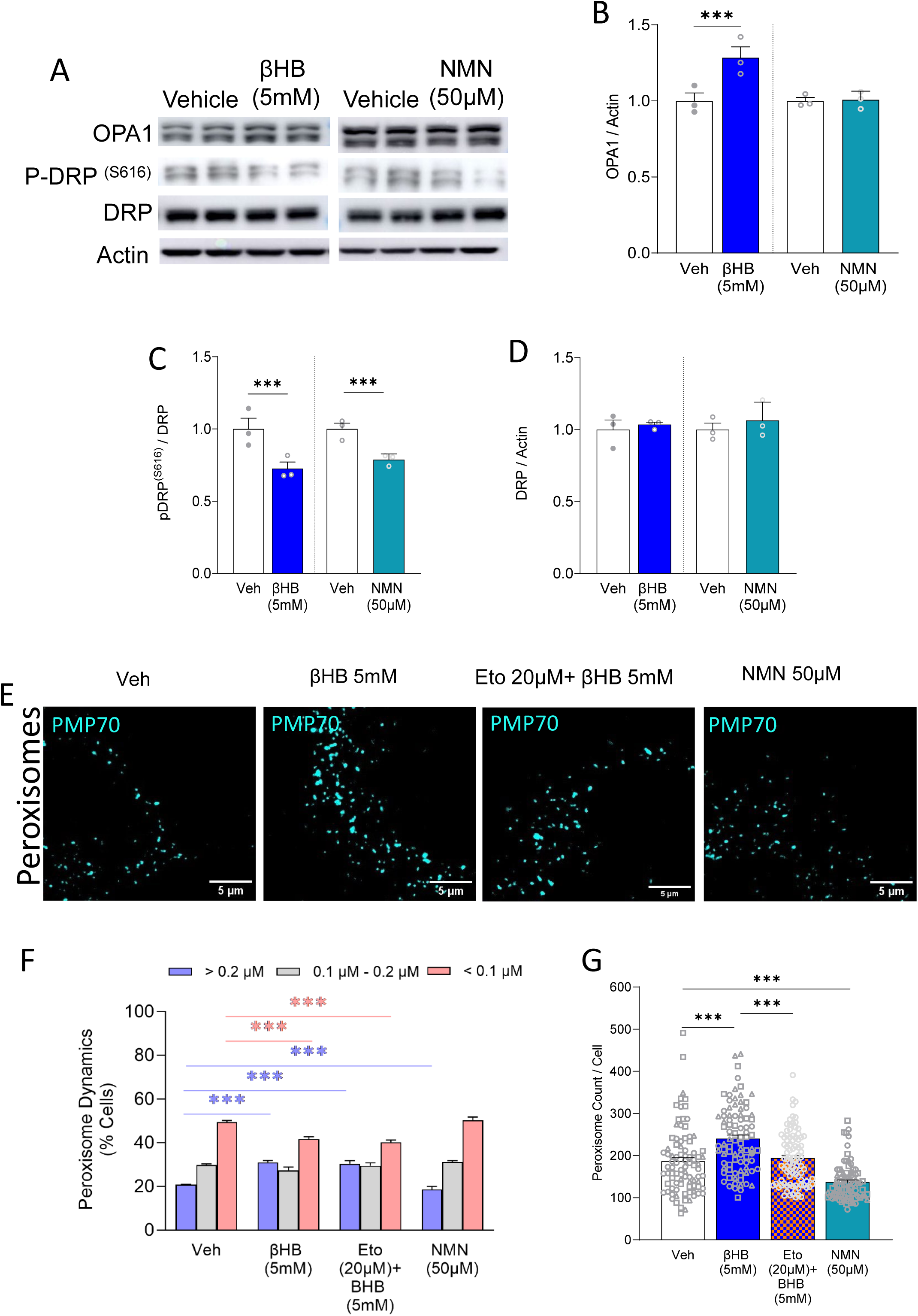
BHB and NMN acts on mitochondrial fusion and fission machinery and peroxisome differently. **(A)** Representative blots are presented for the densitometry analysis for (B) OPA1 and (C) pDRPs616 (D) total DRP(n=3/group). Separately conducted set of experiments are presented using a dotted line under the same graph. (E) Representative confocal images of HeLa cells stained with antibodies against PMP70 for peroxisomes. HeLa cells were immuno-stained after 24h treatment with indicated concentrations of Veh or BHB or Eto+BHB or NMN. (F) Quantification of mean number of peroxisomes per cell were measured between Veh, BHB or Eto+BHB or NMN treated cells. This quantification was performed by Image-J, particle analysis feature, using a standard region of interest (ROI) and measured randomly from all images, with the particle analysis tool. Statistical differences within the individual categories are colour-coded (blue = >0.2 µm, gray = 0.1-0.2 µm, and pink = <0.1 µm) (G) Quantitative analysis of sizes of peroxisomes for control and drug treatment(s) and concentration(s) mentioned. Analysis was done using at least 150 cells per condition, and 3 independent replicates for each treatment condition. Data are shown as mean ± SEM. Data were analyzed using t-test. The significant differences are presented as *** p < 0.001

As peroxisomes share the same fission machinery with mitochondria, but do not undergo fusion with one another, we next examined if ΒHB and NMN also impacted peroxisomal morphology through inhibition of peroxisome fission. Curiously, while ΒHB supplementation increases peroxisome size, NMN supplementation had no effect, suggesting that any effects of ΒHB on peroxisome morphology was independent of NAD^+^ (Fig 2E-F). Supporting this notion, blocking β-oxidation with etoxomir treatment did not impact ΒHB-induced enlargement of peroxisomes. We also noted that ΒHB, but not NMN, induced a marked increase in the number of peroxisomes (Fig 2G), consistent with a role for ΒHB in promoting peroxisome biogenesis independent of its NAD^+^-mediated effects.

Next, we interrogated protein acetylation since increased NAD^+^ promotes protein deacetylation, a post-translational modification linked to changes in mitochondrial morphology (Uddin et al., 2021). Notably, deacetylation of mitochondrial fusion proteins MFN1, MFN2, and OPA1 is reported to promote mitochondrial fusion (Biel et al., 2016; Lee et al., 2014; Samant et al., 2014), while deacetylation of DRP1 is linked to decreased fission (Hu et al., 2020). As expected, we found that ΒHB and NMN both reduce global protein acetylation (Figure 3A and B). More specifically, we observed decreased acetylation of OPA1 and DRP1 (Figure 3A, C and D), but were unable to detect any acetylation of MFN1/ MFN2 proteins.

**Figure 3:**
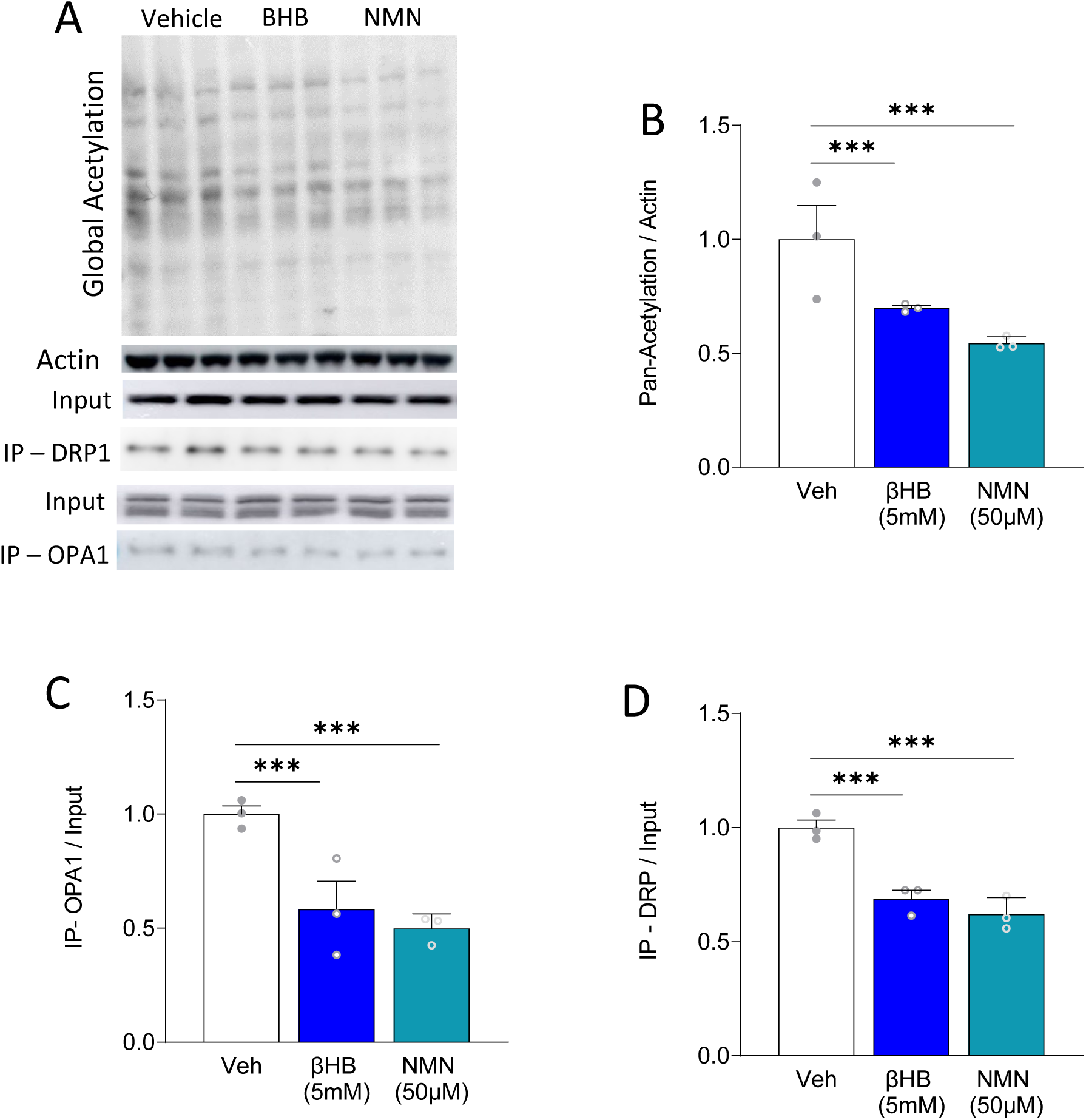
BHB and NMN supplementation in HeLa cell reduces acetylation. **(A)** Representative blots are presented for the densitometry analysis for **(B)** global acetylation in Veh or BHB or NMN treated cells (n=3/group) along with Actin as the loading control; **(C)** acetylated-OPA1 and **(D)** acetylated-DRP1 normalised to input as control (n=3/group). Data are shown as mean ± SEM. Data were analyzed using t-test. The significant differences are presented as *** *p* < 0.001.

### ΒHB and NMN control mitochondrial morphology via sirtuin-based deacetylation activity

To determine if NAD^+^-mediated protein deacetylation is important for promoting mitochondrial elongation induced by ΒHB and NMN, and whether sirtuin deacetylases are involved, we examined the effects of sirtinol, an inhibitor of both Sirt1 and Sirt2 (Mai et al., 2005). Treatment with 50 µM sirtinol increased global acetylation (S4G-H) and caused an accumulation of NAD^+^ (S4I), indicating that the inhibitor was functional. Sirtinol treatment also led to fragmentation of the mitochondrial network (Figure 4A-C), consistent with increased protein acetylation levels correlating to mitochondrial fragmentation. Critically, sirtinol treatment prevented ΒHB and NMN-induced mitochondrial elongation (Figure 4A-C), implicating a role for Sirt1 and/or Sirt2 in ΒHB and NMN-induced mitochondrial elongation.

**Figure 4:**
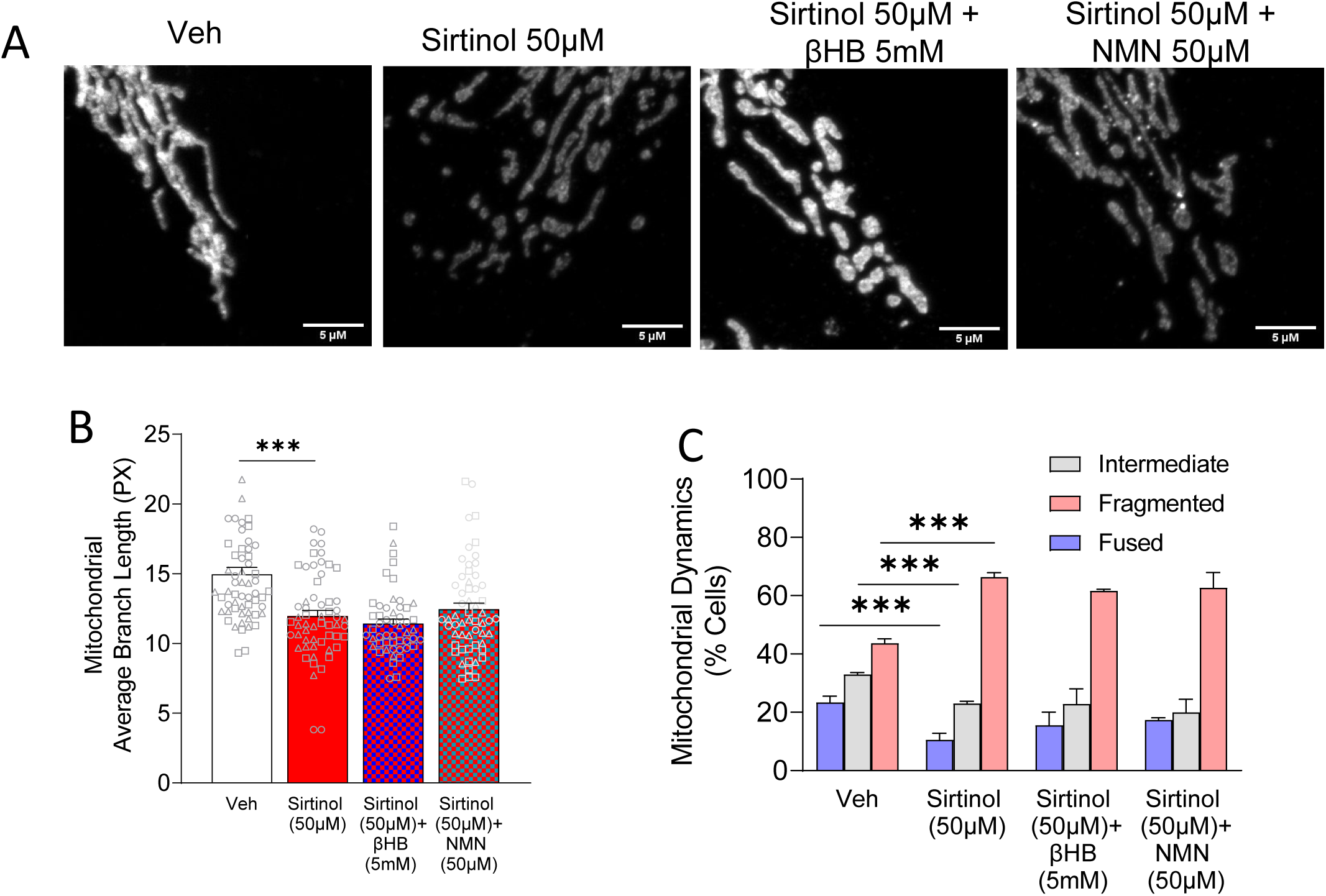
Inhibition of Sirtuin 1 and 2, blocks BHB and NMN aided mitochondrial fusion but not the peroxisome network. **(A)** Representative confocal images of HeLa cells stained with antibodies against TOMM20 for mitochondrial network. HeLa cells were immuno-stained after 24h treatment with indicated concentrations of Veh (Vehicle) or Sirtinol or Sirtinol+BHB or Sirtinol+NMN. (B) Quantification of mean mitochondrial branch length and (C) Qualitative analysis of mitochondrial morphology were measured between Veh or Sirtinol, Sirtinol+BHB or Sirtinol+NMN treated cells. Analysis was done using at least 150 cells per condition, and 3 independent replicates for each treatment condition. (D) Representative confocal images of HeLa cells stained with antibodies against PMP70 for peroxisomes. HeLa cells were immuno-stained after 24h treatment with indicated concentrations of Veh (Vehicle) or Sirtinol or Sirtinol+BHB or Sirtinol+NMN. (E) Quantification of total number of peroxisomes per cell and (F) quantitative analysis of peroxisome size(s) were measured between Veh or Sirtinol, Sirtinol+BHB or Sirtinol+NMN treated cells. Analysis was done using at least 150 cells per condition, and 3 independent replicates for each treatment condition. Data are shown as mean ± SEM. Data were analyzed using t-test. The significant differences are presented as *** p < 0.001.

### Sirt2 is required for NAD^+^ induced mitochondrial elongation via ΒHB

To determine if Sirt1, Sirt2, and/or Sirt3 is required for ΒHB or NMN induced mitochondrial elongation, their expression was knocked down separately via siRNA (Figure 5, S5-7). To control for off-target effects, we used three different siRNA targets for each gene and confirmed knockdown via western blotting (Figure 5D). Upon examining mitochondrial morphology, we observed a similar trend of mitochondrial fragmentation for 2 of the 3 siRNA targets for Sirt1(siRNA#1&2), Sirt2 (siRNA#2&3) and Sirt3 (siRNA#1&3) (S5, S6 and S7), again consistent with the notion that deacetylation promotes mitochondrial elongation. Since the results from Sirt1 siRNA#3, Sirt2 siRNA#1 and Sirt3 siRNA#2 were not consistent (possibly due to off-target effects), they were discarded from our interpretation. Overall, we found that ΒHB-induced mitochondrial elongation was unaffected by Sirt1 and Sirt3 knock-down, but was abolished by Sirt2 knock-down. Meanwhile, knock-down of either Sirt1 or Sirt2 prevented an NMN-induced mitochondrial hyperfusion phenotype, while Sirt3 knock-down had no effect. However, it should be noted that Sirt3 knockdown was less efficient, and that there could still be residual protein activity. Together, our findings indicate that Sirt2 is required for both ΒHB- and NMN-induced mitochondrial elongation via increased NAD^+^ levels. Meanwhile, the fact that ΒHB, but not NMN, can induce mitochondrial elongation in the absence of Sirt1 suggests that ΒHB can also induce mitochondrial elongation via an NAD^+^-independent mechanism, consistent with the fact that ΒHB, but not NMN, increased OPA1 expression.

**Figure 5:**
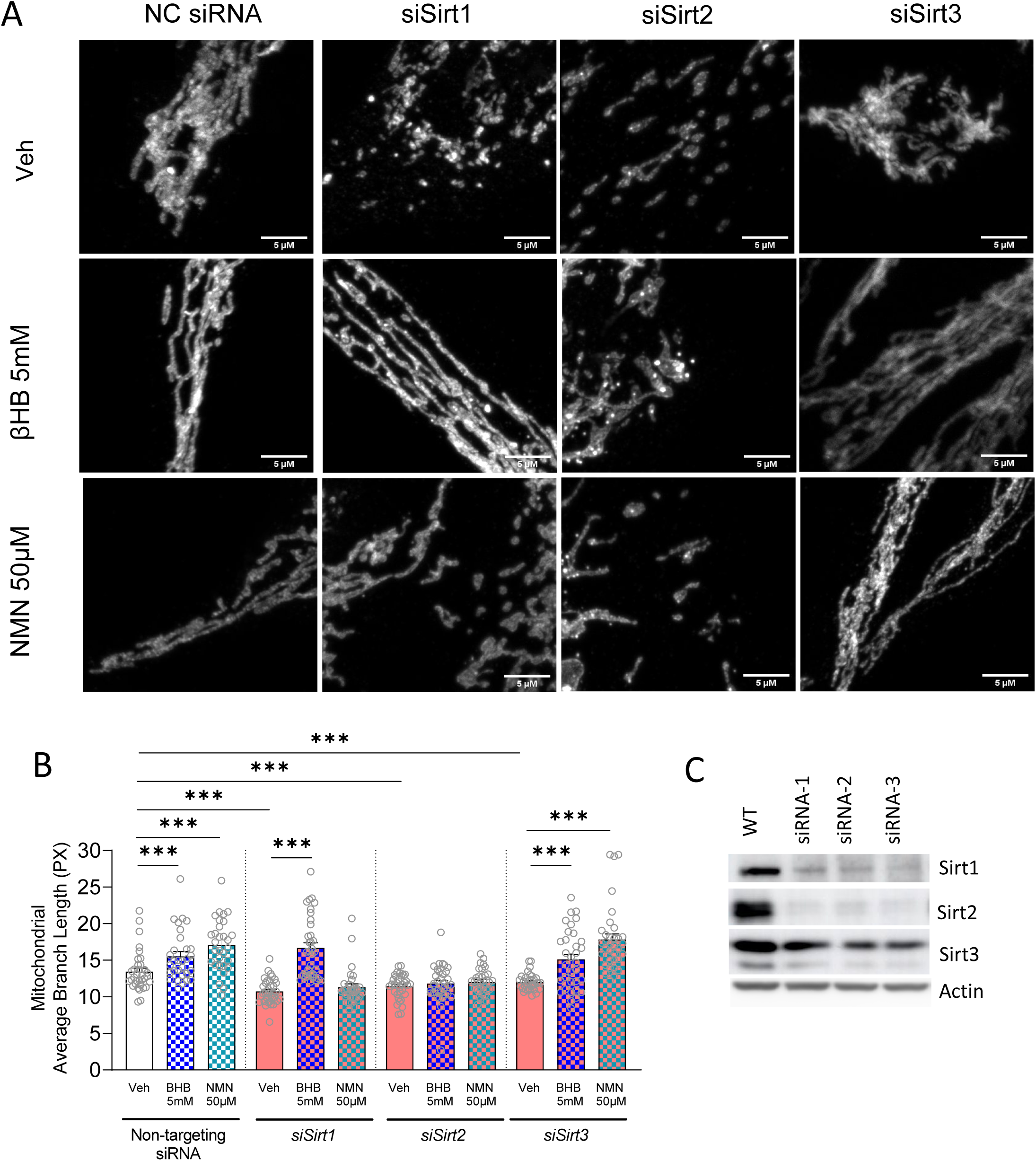
BHB promotes mitochondrial fusion via Sirt2 dependent manner. **(A)** Representative confocal images of HeLa cells stained with antibodies against TOMM20 for mitochondrial network. HeLa cells were immuno-stained 48h post transfection of siRNA targets. 24h post siRNA transfection, with indicated concentrations of BHB or NMN was added for 24h into each siRNA target group. Representative confocal images of HeLa cells stained with antibodies against TOMM20 for mitochondrial network in Sirt1 and Sirt2 KD cells with 24h post treatment of BHB or NMN. Quantification of mean mitochondrial branch length were measured between (B) Veh or BHB or NMN. This quantification was performed by Image-J, MiNA plugin from three independent replicates, using a standard region of interest (ROI) and measured randomly from each image. (C) Qualitative analysis of mitochondrial morphology was performed in Veh, BHB or NMN treated cells via an unbiased visual scoring method. Analysis was done using at least 80 cells per condition, and 2 technical replicates for each treatment condition. Data are shown as mean ± SEM. Data were analyzed using t-test. The significant differences are presented as *** p < 0.001.

### ΒHB and NMN impact behaviours in a *shank3b*^+/−^ zebrafish model of ASD

To examine the efficacy of ΒHB or NMN *in vivo* for the potential treatment of ASD, we studied their effects in *shank3b^+/−^* zebrafish. Wild-type siblings and s*hank3b^+/−^*fish were exposed to vehicle, ΒHB, or NMN over the neurogenic developmental timepoint (10-48 hpf (hour post fertilization); Figure 6A). When fish begin to move at 5 dpf (day post fertilization), locomotor behaviours were examined by measuring total movement in the light and the dark (Figure 6). Specifically, *shank3b^+/−^* fish showed reduced distance travelled in both dark (Figure 6C) and light (Figure 6D); p<0.0001) compared to *shank3b^+/+^* (wild-type) fish exposed to vehicle. Despite a reduced distance travelled, the average speed of *shank3b^+/−^*fish was increased in both light and dark conditions (Fig 6E,F). In *shank3b^+/−^*fish, ΒHB restored distance travelled in light and dark conditions to the levels of wild-type control fish, while there was no difference between the speed of *shank3b^+/−^* and wild-type exposed to BHB. NMN restored distance travelled by *shank3b^+/−^* in light to wild-type levels, while in the dark the distance travelled was similar in both *shank3b^+/−^*and wild-type. The speed of NMN exposed fish was the same in the light and *shank3b^+/−^*fish moved faster in the dark.

**Figure 6:**
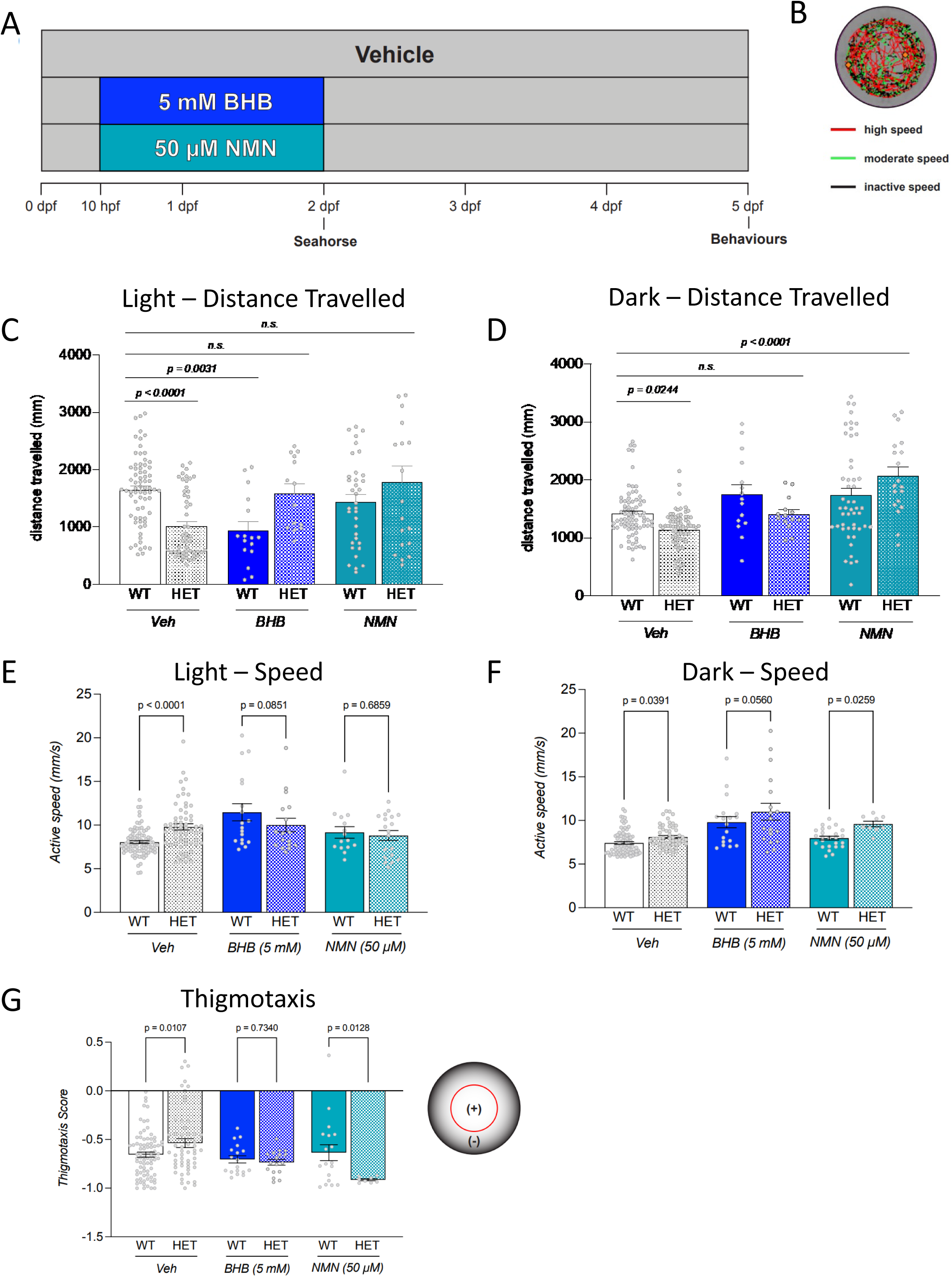
BHB and NMN improve movement behaviours in *shank3b+/−* zebrafish. **A)** Exposure and behavioural paradigm where *shank3b*^+/+^ and *shank3b*^+/−^ zebrafish were exposed to vehicle (Veh; E3), BHB (1 mM), or NMN (50 µM) from 10-48 hpf, and movement monitored at 5 dpf. **B)** Representative movement trace. Analysis of distance travelled (mm) in light **(C)** or dark **(D)** conditions. Analysis of average speed (mm/2) in light **(E)** or dark **(F)** conditions. **G)** Thigmotaxis analysis of zebrafish movement comparing time spent in the center or periphery of the well. Data are shown as mean ± SEM. n=14-71 fish. Data were analyzed using two-way ANOVA with post-hoc Tukey test. Relevant significant differences are presented on the figure.

In addition to standard movement parameters such as distance travelled and speed, we also examined the thigmotaxis, which is the behavioral response to remain close to walls and can be a measure of anxiety. In control conditions, wild-type fish spend more time in the peripheral region near the walls compared to *shank3b^+/−^* fish. However, following exposure to BHB, both wild-type and *shank3b^+/−^* fish spend the same time in the peripheral region. Meanwhile following NMN exposure, *shank3b^+/−^*fish spend more time peripherally than wild-type fish. As such, both BHB and NMN reverse the thigmotaxis differences observed in *shank3b^+/−^* fish.

These findings suggests that either treatment is effective in restoring movement and thigmotaxis to or above the baseline of wild-type, vehicle exposed fish. Unexpectedly, we observed that BHB alone had an effect in wild-type fish under light conditions, significantly decreasing locomotion compared to vehicle-exposed controls (Figure 6C). This decrease was not observed in wild-type fish exposed to NMN, suggesting differences in the mechanism(s) through which ΒHB and NMN elicit their effects. Along these lines, NMN had more dramatic effects on thigmotaxis and standard movement. Nonetheless, these findings indicate that the movement and thigmotaxis phenotypes in *shank3b^+/−^* zebrafish were restored upon developmental exposure to either ΒHB or NMN.

### ΒHB and NMN impact bioenergetics in a *shank3b*^+/−^ zebrafish

We also examined bioenergetics in wild-type and *shank3b^+/−^*fish, and the impact of BHB. At 2 dpf, at the end of the exposure phase, single zebrafish were assessed for bioenergetics using the Seahorse Bioanalyzer to measure oxygen consumption rate (OCR) and extracellular flux (ECAR), basally and in response to various mediators of oxidative phosphorylation (Fig 7). No significant differences were observed in basal respiration upon treatment with oligomycin to block the ATPase or with FCCP treatment to uncouple mitochondria. Of note, upon antimycin A treatment to block complex III and flux through the electron transport chain, *shank3b^+/−^* fish had reduced OCR, indicative of lower non-mitochondrial respiration. However, treatment with BHB rescued this deficit. For ECAR, the only difference between wild-type and *shank3b^+/−^* fish was observed with antimycin A, with *shank3b^+/−^* fish showing a reduction. Upon BHB treatment, no difference between wild-type and *shank3b^+/−^* fish was observed; however, this change appears to be largely due to a decrease in the ECAR in BHB-treated wild-type fish.

**Figure 7:**
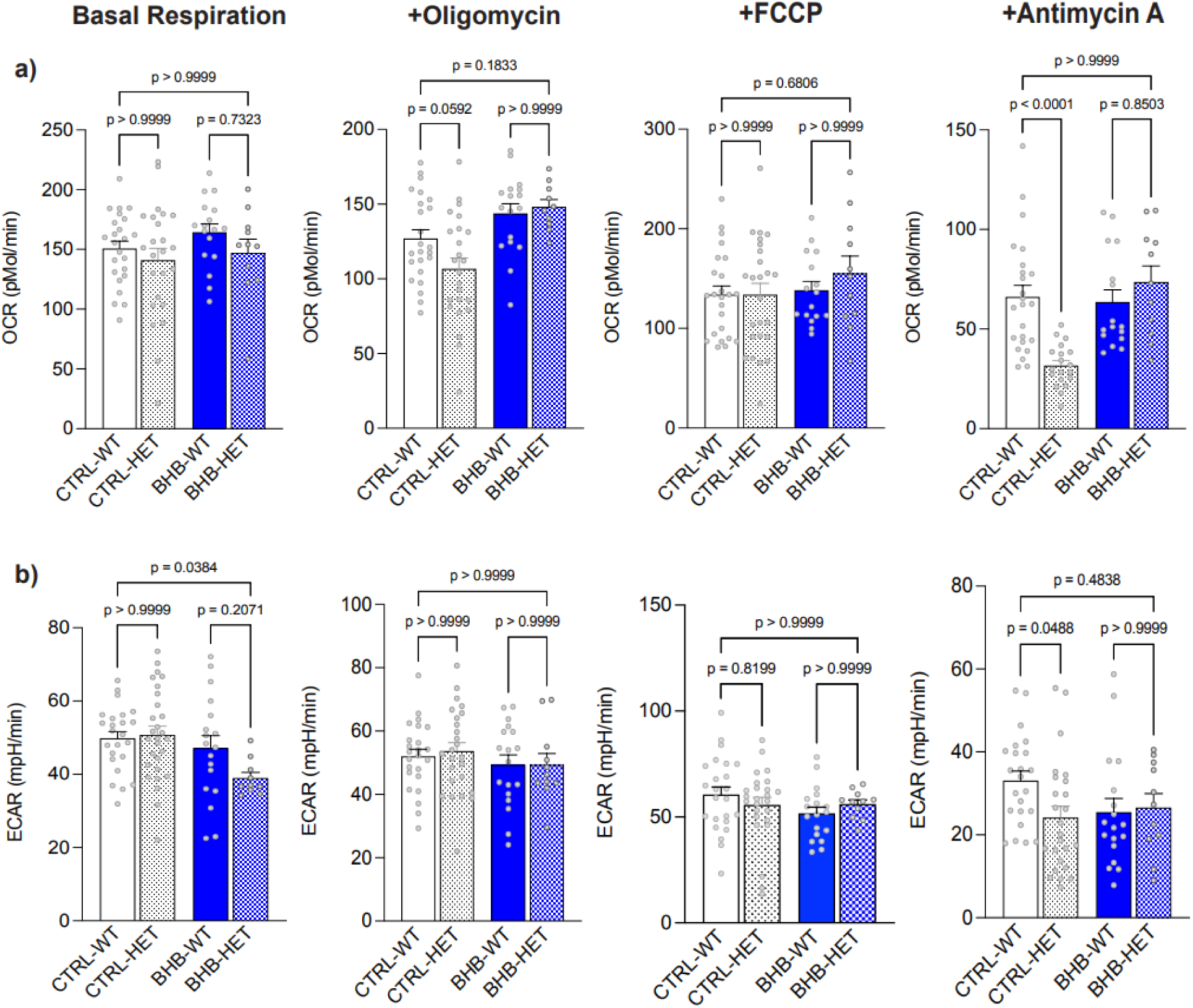
Metabolic effects of BHB and NMN in *shank3+/−* zebrafish. Zebrafish were exposed to vehicle (Veh; E3), BHB (1 mM), or NMN (50 µM) from 10-48 hpf and mitochondrial bioenergetics were measured at 2 dpf. Oxygen consumption rate (OCR; pMol/min) **(A)** and extracellular acidification rate (ECAR; mpH/min) **(B)** were measured basally and following sequential addition of oligomycin, FCCP, and antimycin A as indicated. Data are shown as mean ± SEM. n = 7 per treatment group per run. Data were analyzed using two-way ANOVA with post-hoc Tukey test. Relevant significant differences are presented on the figure.

## Discussion

Our findings provide novel insights into the mechanism by which the KD promotes mitochondrial elongation in HeLa cells and suggest that this pathway might be a promising avenue for improving ASD behaviours *in vivo*. The data herein support a model by which β-oxidation of the KD metabolite ΒHB increases NAD^+^ levels to activate sirtuins, which in turn deacetylate key mitochondrial dynamics proteins to promote mitochondrial hyperfusion. Pathways that modulate mitochondrial morphology often act reciprocally on both fission and fusion, such that mitochondrial hyperfusion is a result of both decreased fission and increased fusion (Sabouny & Shutt, 2020). This situation is true in response to decreased acetylation (Uddin et al., 2021), and our data show this is mirrored by ΒHB and NMN treatments. With respect to increased fusion, both ΒHB and NMN reduced OPA1 acetylation, which promotes mitochondrial fusion (Samant et al., 2014; Sheng & Cristea, 2021). While previous work has shown that deacetylation also activates mitochondrial fusion proteins MFN1 and MFN2 (Lee et al., 2014; Oanh et al., 2017), we were unable to detect acetylation of these proteins to see how they behaved.

Meanwhile, both ΒHB and NMN also reduced DRP1 acetylation, which is known to reduce fission (Hu et al., 2020). Though the mechanism by which deacetylation reduces fission is unknown, it is tempting to speculate that the acetylation status of DRP1 influences the pro-fission phosphorylation at serine-616 of DRP1. This notion is supported by reduced levels of DRP1^P616^ with both ΒHB and NMN treatments in this study, and previous work showing NMN treatment decreased DRP1^P616^ in mice hippocampal tissues (Hu et al., 2020; Klimova et al., 2019), as well as our previous findings of reduced DRP1^P616^ in mouse brains following 2 weeks on the KD (Ahn et al., 2020).

To further investigate the mechanistic pathway by which NAD^+^ mediates deacetylation of proteins mediating mitochondrial dynamics, we interrogated the role of Sirt1 and Sirt2, which both localize to the cytosol where they can act on DRP1, as well as Sirt3, which localizes inside of mitochondria where it can act on OPA1. While Sirt3 knockdown did not affect either ΒHB or NMN-mediated hyperfusion, it is notable that we did not achieve a high reduction in Sirt3 protein levels, and that residual Sirt3 protein could still be active. Meanwhile, knockdown of both Sirt1 and Sirt2 prevented NMN-induced mitochondrial hyperfusion, while only Sirt2 knockdown inhibited ΒHB-induced mitochondrial hyperfusion. This discrepancy could be due to the fact that ΒHB has multiple mechanisms of action beyond the novel pathway we have shown here for protein deacetylation via NAD^+^ and sirtuins (Newman & Verdin, 2014).

In this regard, we also observed other notable differences between ΒHB and NMN treatments. First, while ΒHB increased OPA1 protein levels, NMN did not. This observation is similar to previous work in mouse hippocampal tissues where NMN elevated NAD^+^ levels but did not change OPA1 expression (Klimova et al., 2019). Given that NMN increases NAD^+^ levels, but not total OPA1 protein, we suggest that the elevated OPA1 expression from ΒHB supplementation is independent of NAD^+^ changes. Another difference between ΒHB and NMN was with respect to peroxisome morphology, which was investigated because mitochondria and peroxisomes share the same fission machinery. We propose that the increased peroxisome size and abundance with ΒHB treatment is likely due to the ΒHB induction of peroxisome proliferator-activated receptors (PPARs), rather than increased NAD^+^. This conclusion is based on the observations that etomoxir did not prevent ΒHB-induced peroxisome elongation, and that NMN had no effect on peroxisome size. Moreover, ΒHB-induced increases in peroxisome abundance would be consistent with the known role of PPARs in promoting peroxisome biogenesis and activating β-oxidation (Dreyer et al., 1993; Hess et al., 1965). This alternative pathway likely involves the ability of ΒHB to inhibit histone deacetylases HDAC1, 3 and 4 (Newman & Verdin, 2014; Shimazu et al., 2013), as HDAC3 inhibition activates PPARγ (Galmozzi et al., 2013).

The finding that changes in global acetylation and decreased levels of DRP1^P616^ do not impact peroxisome fission was somewhat unexpected, given the role of DRP1 in the fission of both these organelles. However, this finding is consistent with recent work showing that loss of CLUH did not affect peroxisomal fission despite reducing mitochondrial recruitment of DRP1 and mitochondrial fission (Yang et al., 2022). These unexpected observations suggest that there are discrepancies in how DRP1 regulation impacts both mitochondrial and peroxisomal fission, with DRP1^S616^ affecting mitochondrial fission, but not peroxisome fission

While inhibiting mitochondrial fission or improving mitochondrial fusion alone can improve mitochondrial function in several preclinical disease models (Whitley et al., 2019), it is important to note that we see global decreases in protein deacetylation, not just for mitochondrial fission and fusion proteins. Thus, the benefits of promoting protein deacetylation likely go beyond promoting mitochondrial hyperfusion. Additionally, it should be noted that NAD+ is a critical cofactor for many additional important enzymes that could also contribute to improving mitochondrial function. Consistent with this notion, NAD+ supplementation is emerging as a popular therapeutic strategy in many diseases (Bonkowski & Sinclair, 2016; Gilmour et al., 2020; Uddin et al., 2021).

There is growing evidence that mitochondrial dysfunction, which often correlates with mitochondrial fragmentation, is likely to contribute to ASD (Citrigno et al., 2020; Pecorelli et al., 2020; Rossignol & Frye, 2012; Singh et al., 2020; Thorsen, 2020; Weissman et al., 2008). For example, ASD patient cells (Barone et al., 2021; Frye et al., 2021; Gevezova et al., 2021) and brain organoids (Ilieva et al., 2022) exhibit impaired mitochondrial bioenergetics, while postmortem brains of control and ASD patients have increased expression of mitochondrial fission proteins and decreased expression of mitochondrial fusion proteins (Tang et al., 2013). This correlation is also present in animal models of ASD, as neuronal mitochondria in the BTBR mouse model of ASD have reduced mitochondrial function and are more fragmented (Ahn et al., 2020).

However, despite the benefits of the KD for ASD behaviors in human studies (El-Rashidy et al., 2017; Spilioti et al., 2013) and in animal models (Y. Ahn et al., 2020), we do not as yet fully understand how the KD provides these benefits, as it has many potential modes of action. Nonetheless, the observation that shank3b+/− fish have subtle alterations in bioenergetics, which are restored with BHB exposure, is consistent with a role for mitochondrial dysfunction contributing to ASD. It should be noted that we measured bioenergetics of the whole fish, not the brain specifically; however, our previous studies showed no difference in bioenergetic readouts when measured from whole larval zebrafish or just a dissected head (Ibhazehiebo et al, Brain, 2018), suggesting that the large energy demands of the brain mask any peripheral mitochondrial contributions at this time point.

Our finding that either ΒHB or NMN improve the locomotion and thigmotaxis behaviours of *shank3b^+/−^* fish, begins to provide insight into the mechanisms underlying the benefits of the KD, and is noteworthy for several reasons. First, the observation that BHB alone is beneficial suggests that administration of the full KD may not be necessary. Second, the finding that NMN alone is beneficial suggests that the ability of ΒHB to increase NAD^+^ is important for its benefits. Third, the observation that there are some differences in the effects of BHB and NMN, both at the cellular level and at the behavioural level, suggests that BHB has additional mechanisms of action beyond increasing NAD^+^ that may also impact behaviour. In this regard, it might be interesting to examine the combined effects of BHB and NMN. Finally, if the benefits of ΒHB or NMN in *shank3b^+/−^* fish are translatable to humans, then administration of ΒHB or NMN could be a promising approach that bypasses many of the side effects of the KD. However, while it is tempting to propose BHB or NAD+ boosting treatments for ASD, it should be noted that the locomotion and thigmotaxis assays alone cannot capture the complex behaviours that define ASD. As such, more *in vivo* work with both ΒHB and NMN is required to characterize their effects on additional animal behaviours.

The elucidation of a novel pathway to regulate mitochondrial morphology is an important advancement, given the critical importance of these processes in mediating mitochondrial function, and the increasingly recognized role that mitochondrial dysfunction plays in many disease (Chan, 2020; Whitley et al., 2019). Moreover, restoring the balance between fusion and fission, either by inhibiting fission or by promoting fusion, is beneficial in several neurological disease models. However, there are currently no established direct therapeutic approaches that target mitochondrial fission or fusion in humans. Despite the promise of the KD in this regard, its use may remain somewhat limited due to the risk of undesired side effects and complications. Thus, our findings that ΒHB and NMN treatments can recapitulate some of the beneficial effects of the KD, have promising therapeutic potential.

### Limitations of the study

In the current study, we set out to explore the mechanisms underlying KD-mediated mitochondrial hyperfusion and improvement in ASD behaviours. We identified a mechanistic pathway linking ΒHB, a key metabolite of the KD, to increased NAD^+^ levels and mitochondrial hyperfusion, but there are several limitations to our study. For example, it is important to note that both ΒHB and NAD^+^ have multiple mechanisms of action that likely impact mitochondrial form and function beyond the pathway that we have eluciated. An important future direction would be to investigate the detailed mechanism of how the ketogenic diet (KD) affects NAD+ regulation, including the generation and consumption of NAD+, and the possible involvement of specific enzymes such as iNAMPT, NMNAT1-3, NRK, CD38, and PARPs. In addition, future studies should also explore additional proteins involved in regulating mitochondrial fission and fusion which could also be regulated by acetylation or other mechanisms downstream of BHB and NAD+. Meanwhile, as we did not obtain robust knockdown of Sirt3, we cannot make definitive conclusions about its role in mediating fusion in response to increased NAD^+^. With respect to ASD-related findings, the *shank3^+/−^* fish model is only one ASD model, and it remains unclear if these findings will extend to other ASD models, let alone to human ASD patients. Furthermore, we have only examined movement behaviours, which do not fully recapitulate the full extent of ASD behaviours, and have only examined one timepoint using a single dose for ΒHB and NMN. In this regard, testing additional dosing regimens and behaviours would be important to examine in future studies. Finally, the molecular mechanisms seen in HeLa cells have not been confirmed in zebrafish. Thus, in the future it would be valuable to examine the mitochondrial architecture and in the fish model with and without BHB or NMN treatment, and to determine if these compounds impact mitochondrial morphology through the same molecular mechanisms as in HeLa cells.

## Methods and Materials

### Cell Culture

HeLa cells were grown in DMEM, high-glucose (1g/L) media (Cat#11965092, Gibco, USA) containing l-Glutamine and supplemented with 10% fetal bovine serum (FBS). Cells were seeded at 1 x 10^6^ cells in 10 cm plates and grown at 37 °C and 5% CO_2_ until they were 70-80% confluent (approximately after 48 hrs) before being harvested for analysis or re-seed. At a 50-60% confluency (approximately after 18-24 hrs), cells grown in high-glucose DMEM (1g/L) were treated by the addition of BHB (5 mM) (Cat#H6501; Sigma, USA), NMN (50 μM) (provided by Dr. David Sinclair), etomoxir (20µM) (Cat#509455; Calbiochem, Sigma, USA) or sirtinol (1mM) (Cat#566321; Calbiochem, Sigma, USA), as indicated, for 24 hr. In case of experiments where two drugs were used, both drugs were supplemented in the media together. 24 hr post treatment, cells were harvested by washing with 1x PBS, followed by incubation with 0.25% Trypsin (Corning® Trypsin containing 0.1% EDTA; Fisher Scientific) for 1 min at 37°C and 5% CO_2_. Detached cells were collected in fresh media, followed by centrifugation at 5000 RPM for 5 min, and cell pellets were then stored at −20°C for further experiments. Cells used for immunofluorescence were grown on glass coverslips (no. 1.5 - 12 mm diameter; Electron Microscopy Sciences) that were placed in the same 10cm dishes used for protein analysis.

### Immunofluorescence Staining

Cells on coverslips were fixed in 4% paraformaldehyde for 15 min at 37 °C, permeabilized with 0.1% TritonX100, and blocked with 10% FBS in PBS as described previously (Ahn et al., 2020). Mitochondria were labelled with TOMM20 (Cat#186735; Abcam, USA) primary antibodies were used at 1:1000, diluted in 10% FBS in PBS, and cells were incubated overnight at 4 °C. The appropriate secondary AlexaFluor antibodies were diluted in 10% FBS in PBS at 1:1000, and cells were incubated in the dark at room temperature for 1 hr. Stained cells were then washed in 1X PBS and mounted using Dako mounting media (Cat#S3023, Agilent, USA).

For peroxisome staining, the cells fixed on the coverslips were permeabilized with 0.1% NP-40 dissolved in 1X PBS for 15 minutes at room temperature. This was washed off with PBS and subsequently, 10% FBS was used to block for one hour at room temperature. Following blocking, an overnight incubation was performed to label the peroxisomes with anti-PMP70 (Cat#P0497, Sigma-Aldrich; 1:1000 dilution in blocking solution). The slips were washed three times with PBS before the complementary anti-rabbit secondary antibody was added (Cat#A-11034, Life Technologies; 1:1000 dilution in blocking solution) and incubated for one hour at room temperature. A final washing step with 1X PBS followed and the slips were mounted using the aforementioned mounting media.

### Microscopy and Image Morphology Analysis

An Olympus spinning disc confocal system (Olympus SD OSR, Olympus Corporation, Tokyo, Japan) (UAPON 100XOTIRF/1.49 oil objective) operated by Metamorph software was used to image fixed cells. Mitochondrial network morphology was assessed in a blinded fashion and assigned to 3 categories of mitochondrial morphology (fragmented, intermediate, and fused). A representative image under each category is shown in S3A. At least 80-150 cells were counted per condition, and the analyses were performed on 3 independent replicates and 2 independent technical replicates for knockdown study for each treatment condition. Mitochondrial branch length was quantified using the Image-J, MiNA (Valente et al., 2017) plug-in from three independent replicates, using a standard region of interest (ROI) and measured randomly from each image.

For peroxisome analysis of the various treatments, Z-projections were obtained, and images adjusted for threshold, the threshold was set to remove any background signals using Image J. Z projections were generated for average intensity using Image J. The Image J particle analysis tool was used to measure peroxisome size, and counts.

For peroxisome size analysis, peroxisomes from 200 cells were analyzed and the sizes binned as follows: elongated (0.2-1) µm^2^, intermediate (0.1-0.2) µm^2^ or round (0.01-0.1) µm^2^. Meanwhile, the mean number of peroxisomes was calculated per cell. Sample size is 88 cells in total, with 2 biological replicates (44 cells/replicate).

### Oxygen Consumption

For each well in flat bottom 96 well plate (MaxBioChem), 30,000 HeLa cells were seeded in DMEM culture medium to and allowed to attach overnight. The next day, new media with 5 mM BHB (Cat#H6501; Sigma, USA) or vehicle was added. 24 hours later the oxygen consumption rate (OCR) was measured using the Resipher platform (Lucid Scientific, Atlanta, GA, United States) (Triolo & Khacho, 2024). Real-time continuous OCR measurements were monitored in the incubator. Finally, the OCR was normalized to total protein content estimated using the Pierce™ BCA assay (Cat#A555865; ThermoFisher, USA).

### Western blots

Frozen cell lysates were homogenized using RIPA Lysis and Extraction Buffer (Cat#89900, Thermo Fisher, USA) and protease inhibitors cocktail (Cat#10190-060, VWR, USA), on ice for 1hr. Protein content was measured using the Pierce™ BCA Protein Assay Kit (Cat#A555865; ThermoFisher, USA). Depending on the target, 30–40-μg total protein was loaded onto SDS-PAGE for western blotting.

Polyvinylidene fluoride (PVDF) membranes were blocked with 5% nonfat milk in Tris-buffered saline solution containing 0.1% Tween 20 (TBS-T) for 1 h. After blocking, the blots were incubated with primary antibody overnight at 4°C. After washing with TBS-T, the blots were incubated with the secondary antibody for 1 h at room temperature. After washing with TBS-T, blots were visualized using an SuperSignal™ West Femto Maximum Sensitivity Substrate (Cat#34095; Fisher Scientific, USA). Bands were detected and quantified using a ChemiDOC MP gel imaging system (Bio-Rad). The following antibodies were used: Actin (Cell Signaling; 1:10,000); DRP1 (BD Biosciences, San Jose, CA, USA; 1:1000); pDRP1^S616^ at (Cell Signaling, Danvers, MA, USA; 1:1000); OPA1 (BD Biosciences; 1:1000); MFN1 (Cat# 3455S, Cell Signaling, USA; 1:1000); MFN2 (Cat# M03, Abnova, USA; 1:1000); MFF (Cat#, Cell Signaling, USA; 1:1000); and DRP1 S637 (Cat# 4867SS, Cell Signaling, USA; 1:1000). The expression levels of proteins were normalized to actin for cell lysate, and the ratio of the phosphorylated and total forms was computed using the normalized relative total expression levels. Original blots that were used to crop representative are shown in S8A-F

### Immunoprecipitation

Freshly centrifuged cell pallet from the cell lysate collected after the 24hr treatment (as described in cell culture section) from a 10 cm dish. 250 µl of IP homogenization buffer was used to extract proteins. A custom homogenized buffer was prepared for IP sample homogenization, 150mM NaCL (Cat#BP358212; Fisher Scientific, USA), 50mM Tris-HCL at 7.5 pH (Cat#97063-756; VWR, USA), 5mM EDTA at 8 pH (Cat#E3889; Sigma Adrich, USA), 0.5% NP-40 (Cat#00000; Company), and 1% TritonX (Cat#X100; Sigma Adrich, USA); with freshly added Trichostatin A 1 μM (Cat#A8183; ApexBIO, USA), Nicotinamide 5mM (Cat#N0636; Sigma Adrich, USA), 10mM Sodium Butyrate (Cat#303410; Sigma Adrich, USA) and Protease inhibitor cocktail 1:100 (Cat#M250; Amresco, USA). After 1 hr of incubation, samples were centrifugated at 10000 RPM. Supernatant was diluted accordingly to prepare same amount of protein (350-400ug) and volume (200 µl) followed by protein quantification.

Preclearing and washing cell lysate were performed using 20 µl A/G PLUS-Agarose beads per 200 µl sample (Cat#SC2003; Santa Cruz Biotechnology, USA) for 4 hrs followed by centrifugation at 15,000g for 10 minutes at 4°c. 20 µl of collected supernatant was kept separately to use as input. The leftover supernatant was then incubated with 1.5 µL anti-acetyl-lysine antibody (Cat#9441S; Cell signalling, USA) overnight. 60-70 µl (depending on the protein concentration used in the beginning) A/G PLUS-Agarose beads were added to the antibody incubated samples. Followed by a 4hr rotation and incubation at 4°C, samples were centrifuged at 5,000g for 5 minutes. Supernatant was discarded and the beads pallet was then washed 3 times with the IP homogenization or extraction buffer with all the components. Washing was performed at at 5,000g for 5 minutes each time. Pallets were then heated at 95°C for 5 min with Laemmli buffer (Cat#1610747; Biorad, USA) and centrifuged at 15,000g for 10 minutes to collect the supernatant. This supernatant was used to perform immunoblotting. Input was prepared checked in a separate gel but same day and setup. Also, a pooled samples (10 µl/sample/group) from all groups ran to show as a positive control, 1 sample input /group was used to run together with IP sample to show the differences. IP sample successfully showed heavy chain and light chain produced due to the A/G bead’s reaction. Original blots that were used to crop representative are shown in S8A-F.

### NAD assay

NAD^+^ and NADH were measured as described previously (Zhu & Rand, 2012). Briefly, samples were homogenized in extraction buffer (10 mmol/l Tris/HCl, 0.5% Triton X-100, 10 mmol/l Nicotinamide, pH 7.4) and then centrifuged (12,000 × g for 5 min at 4◦C), after which an aliquot of supernatant was taken for protein quantification. After subsequent extractions with phenol: chloroform: isoamyl alcohol (25:24:1) and chloroform, the supernatant was separated in two aliquots to measure total NAD or NAD+ using a Bio-Rad Imark microplate reader. Data are presented as pmol of NAD^+^ or NADH per mg of protein.

### siRNA knockdow

TriFECTa RNAi Kits (Integrated DNA Technology, USA) were obtained for SIRT1 (hs.Ri.SIRT1.13), SIRT2 (hs.Ri.SIRT2.13) and Sirt3 (hs.Ri.SIRT3.13). Both a transfection control (TYE563 DsiRNA) (Cat#51-01-20-19), and a negative control DsiRNA (a nontargeting DsiRNA that has no known targets in human, mouse, or rat) (Cat#51-01-14-03), were provided with the kit. 24hrs after siRNA transfection, vehicle, ΒHB or NMN treatments were initiated, and carried out an additional 24hr. Lipofectamine™ RNAiMAX (Cat#13778075, Thermo Fisher Scientific, USA) was used as transfection reagent. Cells were platted at 0.5×10^5^ density in a 6 well plate. After 12 hrs of seeding, RNAiMAX (9µl/well) and siRNA (100pmol/well) was diluted in Opti-MEM medium (Cat#31985062, Thermo Fisher Scientific, USA). The mixture of RNAiMAX and siRNA was then added to the well for 24hr. Cells were then collected for western analysis and immunfluorescence to analyse mitochondrial morphology as above.

### Zebrafish husbandry, behaviour and genotyping

Zebrafish heterozygous for a *shank3b* mutation (*shank3b^+/−^*) were placed in breeding tanks the afternoon prior to day 0 of the experiment. Males and females were separated by a removable insert, which was removed the morning of the next day to allow breeding. Embryos were consequently collected in petri dishes at a density of ∼50 embryos/plate and maintained in E3 medium (60X E3 and methylene blue, diluted in milli Q water) at 28°C. At 10 hours-post fertilization (hpf), zebrafish were either exposed to vehicle (E3 medium), BHB (5 mM), or NMN (50 µM). At 48 hpf, all exposure conditions were washed out thrice and replaced by E3 medium. Zebrafish underwent behavioural assays at 5 days-post fertilization (dpf). Briefly, zebrafish were placed in 48-well plates at a density of 1 fish/well. Locomotion, including distance travelled and speed, was analyzed in each of 15-minutes light and 15-minutes dark via tracking assay using the ZebraBox**®** recording chamber (Viewpoint Life Sciences, Montreal, QC). Thigmotaxis was measured by comparing the time spent (light conditions) in the inside region of the well versus the perimeter of the well. The thigmotaxis score ranged from −1 to +1 and was calculated with the following formula: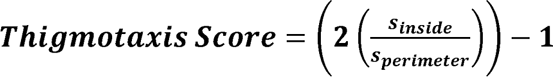. After behavioural assays were completed, zebrafish were genotyped by PCR amplification of the *shank3b* gene with subsequent digestion with T7 endonuclease I (M0302; New England BioLabs). Genotypes were verified on a 2% agarose gel. Mismatched duplexes (seen as three bands on a gel) indicated a heterozygous genotype, whereas single banded samples required further DNA sequencing to indicate a wild type versus homozygous genotype.

### Zebrafish oxygen consumption

2 dpf larval zebrafish were used to measure mitochondrial bioenergetics upon 5 mM BHB exposure from 10 hpf to 2 dpf. Oxygen consumption rates (OCR; pMol/min) and extracellular acidification rates (ECAR; mpH/min) were measured using a XF24 Extracellular Analyzer (Seahorse Biosciences, Billerica, MA). Single zebrafish (n = 7 per treatment group per run) were seeded in an islet microplate, protected by a mesh islet capture screen. To assess zebrafish bioenergetics, we followed a previously published methodology (Stackley et al., 2011). In brief, OCR and ECAR were measured across 27 cycles (2-minutes mixing, 1-minute wait, 2-minute measurement) at basal rates and after exposure to dilutions of 100 µM oligomycin (injected at cycle 7), 37.5 µM carbonyl cyanide p-trifluoromethoxyphenylhydrazone (FCCP; injected at cycle 15), and 30 µM antimycin A (injected at cycle 21). Final measurements were taken as the average of the last 4 cycles for basal respiration, last 3 cycles for oligomycin and antimycin A injections, and last 5 cycles for FCCP injections.

### Statistics

Statistical analysis was performed using GraphPad Prism 9 (Version 9.4.1) software (La Jolla, CA, USA). All data are presented as mean ± SEM. Mechanistic data were analyzed using unpaired t-tests to assess significant changes between two groups (e.g., vehicle vs. NMN or vehicle vs. BHB). For the zebrafish assay, data involving multiple factors were analyzed using two-way ANOVA followed by Tukey’s post-hoc test. Normality was assessed using the Shapiro-Wilk test for all datasets, and no violations of normality were observed. Significant differences are presented as *p < 0.05, **p < 0.01, and **p < 0.001.

## Authors Contribution

GMU: designed the study, performed experiments, analysed data, coordinated experiments with other collaborators, generated figures, and wrote and edited the manuscript. RL: designed and performed fish behavioural assays (including breeding) and Seahorse assays, analysed data, and contributed to writing and editing. MZ: performed the peroxisome assay and data analysis. AJ: performed cellular oxygen consumption assays IAK: provided technical support for microscopy and siRNA experiments. JMR: obtained funding, designed the study, and edited manuscript. DMK: designed the fish experiments, and edited the manuscript. TES: obtained funding, designed and coordinated the study, and wrote and edited the manuscript as the corresponding author.

## Acknowledgements

We thank Dr David A. Sinclair (Professor at the Department of Genetics and Co-Director of the Paul F. Glenn Center for Biology of Aging Research at Harvard Medical School) for kindly providing the NMN.

## Ethics Approval

All fish protocols were approved by the University of Calgary Institutional Animal Care and Use Committee, which conform with the Guide for the Care and Use of Laboratory Animals published by the United States National Institutes of Health (eighth edition; revised 2011) and the guidelines of the Canadian Council on Animal Care.

## Sources of Funding

This study was supported by funds from the Scottish Rites Charitable Foundation (TES, JMR), and the Alberta Children’s Hospital Research Foundation Neurodevelopment Disorder Program (JMR, TES). MZ was supported by Owerko Center, University of Calgary. GMU was supported by an Alberta Children’s Hospital Research Foundation - Cumming School of Medicine postdoctoral fellowship. The funders had no role in study design, data collection and analysis, decision to publish, or preparation of the manuscript.

## Conflict of interest

The authors have declared that no conflict of interest exists

## Supporting Information

**S1 Figure. BHB promotes mitochondrial elongation.** Qualitative analyses of mitochondrial network morphology of HeLa cells following BHB treatment. Morphology is characterized according to the appearance of the mitochondrial network as fused, intermediate, or fragmented, via an unbiased scoring method. Analyses was performed from a total of 150 cells per treatment, with 3 independent replicates for each treatment. Bars indicate mean +/− SD.

**S2 Figure. BHB promotes mitochondrial respiration.** HeLa cells were pre-treated with BHB or vehicle control for 24 hrs and the basal oxygen consumption rate OCR (fmol/mm^2^/s/mg protein) acquired via the Resipher platform. **A)** Basal OCR trace acquired over 10 hrs. Error bars show mean +/− SD. n = 4/group **B)** Basal OCR at 25 hour and 33 hour time points.). Bars show median and points represent replicates. Data were analyzed via unpaired t-test; * p ≤ 0.05; ** p ≤ 0.01; *** p ≤ 0.001; ns, not significant.

**S3 Figure. Impact of BHB and NMN on cellular NAD+ levels and mitochondrial dynamics proteins. (A)** Representative images for the visual scoring of mitochondrial morphology in HeLa cells. **(B)** NAD^+^ and **(C)** Total NAD^+^+NADH level was measured 24h post treatment of Veh or BHB or NMN n HeLa cells. **(D)** Representative blots are presented for the densitometry analysis for mitochondrial fusion and fission related protein expression of **(E)** MFN1, **(F)** MFN2, **(G)** MFF in Veh, and BHB or NMN treated HeLa cells (n=3/group). Data are shown as mean ± SEM. Data were analyzed using t-test. The significant differences are presented as *** *p* < 0.001.

**S4 Figure. β-oxidation of BHB is necessary to promote mitochondrial elongation. (A)** Representative confocal images of HeLa cells stained with antibodies against TOMM20 for mitochondrial network. HeLa cells were immuno-stained after 24h treatment with indicated concentrations of Veh or BHB or Eto or Eto+BHB. **(B)** Quantification of mitochondrial average branch length were measured between Veh or BHB or Eto or Eto+BHB treated cells. This quantification was performed by Image-J, MiNA plugin from three independent replicates, using a standard region of interest (ROI) and measured randomly from each image. **(C)** Qualitative analysis of mitochondrial morphology was performed in Veh or BHB or Eto or Eto+BHB treated cells via an unbiased visual scoring method. Analysis was done using at least 150 cells per condition, and 3 independent replicates for each treatment condition. **(D, E)** NAD^+^ and total NAD^+^+NADH level was measured 24h post treatment of Veh, BHB, Eto or Eto+BHB. **(F, G)** Representative blots of global acetylation in Veh, or sirtinol treated cells for 24h (n=3/group) along with Actin as the loading control and their densitometric analysis. **(H, I**) NAD^+^ and total NAD^+^+NADH level was measured 24h post treatment of Veh, Sirtinol, Sirtinol+BHB or Sirtinol+NMN in HeLa cells respectively. Data are shown as mean ± SEM. Data were analyzed using t-test. The significant differences are presented as *** *p* < 0.001.

**S5 Figure. SIRT1 knockdown prevents NMN induced mitochondrial elongation. (A)** Representative confocal images of HeLa cells stained with antibodies against TOMM20 for mitochondrial network. **(A)** The HeLa cells were immuno-stained 48h post transfection of siSirt1 (target1, target 2 and target 3). 24h post siRNA transfection, with indicated concentrations of BHB or NMN was added for 24h into each siRNA target group. **(B)** Quantification of mitochondrial average branch length were measured between the indicated groups in each siRNA targets. This quantification was performed by Image-J, MiNA plugin from three independent replicates, using a standard region of interest (ROI) and measured randomly from each image. **(C)** Qualitative quantification of mitochondrial morphology was performed between the indicated groups in each siRNA targets via an unbiased visual scoring method. Analysis was done using at least 80 cells per condition, and 2 technical replicates for each treatment condition. Data are shown as mean ± SEM. Data were analyzed using t-test. The significant differences are presented as *** *p* < 0.001.

**S6 Figure. SIRT2 knockdown prevents BHB and NMN induced mitochondrial elongation. (A)** Representative confocal images of HeLa cells stained with antibodies against TOMM20 for mitochondrial network. **(A)** The HeLa cells were immuno-stained 48h post transfection of siSirt2 (target1, target 2 and target 3). 24h post siRNA transfection, with indicated concentrations of BHB or NMN was added for 24h into each siRNA target group. **(B)** Quantification of mitochondrial average branch length were measured between the indicated groups in each siRNA targets. This quantification was performed by Image-J, MiNA plugin from three independent replicates, using a standard region of interest (ROI) and measured randomly from each image. **(C)** Qualitative quantification of mitochondrial morphology was performed between the indicated groups in each siRNA targets via an unbiased visual scoring method. Analysis was done using at least 80 cells per condition, and 2 technical replicates for each treatment condition. Data are shown as mean ± SEM. Data were analyzed using t-test. The significant differences are presented as *** *p* < 0.001.

**S7 Figure. Effects of SIRT3 knockdown on BHB and NMN induced mitochondrial elongation. (A)** Representative confocal images of HeLa cells stained with antibodies against TOMM20 for mitochondrial network. **(A)** The HeLa cells were immuno-stained 48h post transfection of siSirt3 (target1, target 2 and target 3). 24h post siRNA transfection, with indicated concentrations of BHB or NMN was added for 24h into each siRNA target group. **(B)** Quantification of mitochondrial average branch length were measured between the indicated groups in each siRNA targets. This quantification was performed by Image-J, MiNA plugin from three independent replicates, using a standard region of interest (ROI) and measured randomly from each image. **(C)** Qualitative quantification of mitochondrial morphology was performed between the indicated groups in each siRNA targets via an unbiased visual scoring method. Analysis was done using at least 80 cells per condition, and 2 technical replicates for each treatment condition. Data are shown as mean ± SEM. Data were analyzed using t-test. The significant differences are presented as *** *p* < 0.001.

**S8 Figure. Uncropped immunoblots. (A-F)** Original blots presented that was used to present the representative bands in analysis and figures.

## Notes

### Competing Interest Statement

The authors have declared no competing interest.

### Summary of Updates

Updated version of the manuscript as part of revisions for a journal review.

## References

Ahn, Sabouny, R., Villa, B. R., Yee, N. C., Mychasiuk, R., Uddin, G. M.,…Shutt, T. E. (2020). Aberrant Mitochondrial Morphology and Function in the BTBR Mouse Model of Autism Is Improved by Two Weeks of Ketogenic Diet. Int J Mol Sci, 21(9). 10.3390/ijms21093266

Ahn, Y., Narous, M., Tobias, R., Rho, J. M., & Mychasiuk, R. (2014). The ketogenic diet modifies social and metabolic alterations identified in the prenatal valproic acid model of autism spectrum disorder. Dev Neurosci, 36(5), 371–380. 10.1159/000362645

Ahn, Y., Sabouny, R., Villa, B. R., Yee, N. C., Mychasiuk, R., Uddin, G. M.,…Shutt, T. E. (2020). Aberrant mitochondrial morphology and function in the BTBR mouse model of autism is improved by two weeks of ketogenic diet. International journal of molecular sciences, 21(9), 3266.

Baker, N., Patel, J., & Khacho, M. (2019). Linking mitochondrial dynamics, cristae remodeling and supercomplex formation: how mitochondrial structure can regulate bioenergetics. Mitochondrion, 49, 259–268.

Barone, R., Bastin, J., Djouadi, F., Singh, I., Karim, M. A., Ammanamanchi, A.,…Frye, R. E. (2021). Mitochondrial Fatty Acid beta-Oxidation and Resveratrol Effect in Fibroblasts from Patients with Autism Spectrum Disorder. J Pers Med, 11(6). 10.3390/jpm11060510

Biel, T. G., Lee, S., Flores-Toro, J. A., Dean, J. W., Go, K. L., Lee, M. H.,…Kim, J. S. (2016). Sirtuin 1 suppresses mitochondrial dysfunction of ischemic mouse livers in a mitofusin 2-dependent manner. Cell Death Differ, 23(2), 279–290. 10.1038/cdd.2015.96

Bonkowski, M. S., & Sinclair, D. A. (2016). Slowing ageing by design: the rise of NAD(+) and sirtuin-activating compounds. Nat Rev Mol Cell Biol, 17(11), 679–690. 10.1038/nrm.2016.93

Chan, D. C. (2020). Mitochondrial dynamics and its involvement in disease. Annual review of pathology: mechanisms of disease, 15, 235–259.

Citrigno, L., Muglia, M., Qualtieri, A., Spadafora, P., Cavalcanti, F., Pioggia, G., & Cerasa, A. (2020). The Mitochondrial Dysfunction Hypothesis in Autism Spectrum Disorders: Current Status and Future Perspectives. Int J Mol Sci, 21(16). 10.3390/ijms21165785

Covarrubias, A. J., Perrone, R., Grozio, A., & Verdin, E. (2021). NAD(+) metabolism and its roles in cellular processes during ageing. Nat Rev Mol Cell Biol, 22(2), 119–141. 10.1038/s41580-020-00313-x

Dreyer, C., Keller, H., Mahfoudi, A., Laudet, V., Krey, G., & Wahli, W. (1993). Positive regulation of the peroxisomal beta-oxidation pathway by fatty acids through activation of peroxisome proliferator-activated receptors (PPAR). Biol Cell, 77(1), 67–76. 10.1016/s0248-4900(05)80176-5

Durand, C. M., Betancur, C., Boeckers, T. M., Bockmann, J., Chaste, P., Fauchereau, F.,…Bourgeron, T. (2007). Mutations in the gene encoding the synaptic scaffolding protein SHANK3 are associated with autism spectrum disorders. Nat Genet, 39(1), 25–27. 10.1038/ng1933

El-Rashidy, O., El-Baz, F., El-Gendy, Y., Khalaf, R., Reda, D., & Saad, K. (2017). Ketogenic diet versus gluten free casein free diet in autistic children: a case-control study. Metab Brain Dis, 32(6), 1935–1941. 10.1007/s11011-017-0088-z

Elamin, M., Ruskin, D. N., Masino, S. A., & Sacchetti, P. (2017). Ketone-Based Metabolic Therapy: Is Increased NAD(+) a Primary Mechanism? Front Mol Neurosci, 10, 377. 10.3389/fnmol.2017.00377

Evangeliou, A., Vlachonikolis, I., Mihailidou, H., Spilioti, M., Skarpalezou, A., Makaronas, N.,…Smeitink, J. (2003). Application of a ketogenic diet in children with autistic behavior: pilot study. J Child Neurol, 18(2), 113–118. 10.1177/08830738030180020501

Fang, E. F., Hou, Y., Palikaras, K., Adriaanse, B. A., Kerr, J. S., Yang, B.,…Bohr, V. A. (2019). Mitophagy inhibits amyloid-β and tau pathology and reverses cognitive deficits in models of Alzheimer’s disease. Nat Neurosci, 22(3), 401–412. 10.1038/s41593-018-0332-9

Fang, E. F., Scheibye-Knudsen, M., Brace, L. E., Kassahun, H., SenGupta, T., Nilsen, H.,…Bohr, V. A. (2014). Defective mitophagy in XPA via PARP-1 hyperactivation and NAD(+)/SIRT1 reduction. Cell, 157(4), 882–896. 10.1016/j.cell.2014.03.026

Frye, R. E., Lionnard, L., Singh, I., Karim, M. A., Chajra, H., Frechet, M.,…Aouacheria, A. (2021). Mitochondrial morphology is associated with respiratory chain uncoupling in autism spectrum disorder. Transl Psychiatry, 11(1), 527. 10.1038/s41398-021-01647-6

Galmozzi, A., Mitro, N., Ferrari, A., Gers, E., Gilardi, F., Godio, C.,…Crestani, M. (2013). Inhibition of class I histone deacetylases unveils a mitochondrial signature and enhances oxidative metabolism in skeletal muscle and adipose tissue. Diabetes, 62(3), 732–742. 10.2337/db12-0548

Gevezova, M., Minchev, D., Pacheva, I., Sbirkov, Y., Yordanova, R., Timova, E.,…Sarafian, V. (2021). Cellular Bioenergetic and Metabolic Changes in Patients with Autism Spectrum Disorder. Curr Top Med Chem, 21(11), 985–994. 10.2174/1568026621666210521142131

Gilmour, B. C., Gudmundsrud, R., Frank, J., Hov, A., Lautrup, S., Aman, Y.,…Fang, E. F. (2020). Targeting NAD(+) in translational research to relieve diseases and conditions of metabolic stress and ageing. Mech Ageing Dev, 186, 111208. 10.1016/j.mad.2020.111208

Glancy, B., Kim, Y., Katti, P., & Willingham, T. B. (2020). The functional impact of mitochondrial structure across subcellular scales. Frontiers in physiology, 11.

Gogou, M., & Kolios, G. (2018). Are therapeutic diets an emerging additional choice in autism spectrum disorder management? World J Pediatr, 14(3), 215–223. 10.1007/s12519-018-0164-4

Hess, R., Staubli, W., & Riess, W. (1965). Nature of the hepatomegalic effect produced by ethyl-chlorophenoxy-isobutyrate in the rat. Nature, 208(5013), 856–858. 10.1038/208856a0

Hu, Q., Zhang, H., Gutierrez Cortes, N., Wu, D., Wang, P., Zhang, J.,…Wang, W. (2020). Increased Drp1 Acetylation by Lipid Overload Induces Cardiomyocyte Death and Heart Dysfunction. Circ Res, 126(4), 456–470. 10.1161/CIRCRESAHA.119.315252

Ibhazehiebo K., Gavrilovici C., de la Hoz C.L., Ma S.C., Rehak R., Kaushik G., Meza Santoscoy P.L., Scott L., Nath N., Kim D.Y., Rho J.M., Kurrasch D.M. (2018). A novel metabolism-based phenotypic drug discovery platform in zebrafish uncovers HDACs 1 and 3 as a potential combined anti-seizure drug target. Brain. 10.1093/brain/awx364

Ilieva, M., Aldana, B. I., Vinten, K. T., Hohmann, S., Woofenden, T. W., Lukjanska, R.,…Michel, T. M. (2022). Proteomic phenotype of cerebral organoids derived from autism spectrum disorder patients reveal disrupted energy metabolism, cellular components, and biological processes. Mol Psychiatry. 10.1038/s41380-022-01627-2

Kalueff, A. V., Stewart, A. M., & Gerlai, R. (2014). Zebrafish as an emerging model for studying complex brain disorders. Trends Pharmacol Sci, 35(2), 63–75. 10.1016/j.tips.2013.12.002

Kang, H. C., Chung, D. E., Kim, D. W., & Kim, H. D. (2004). Early- and late-onset complications of the ketogenic diet for intractable epilepsy. Epilepsia, 45(9), 1116–1123. 10.1111/j.0013-9580.2004.10004.x

Kartawy, M., Khaliulin, I., & Amal, H. (2021). Systems Biology Reveals S-Nitrosylation-Dependent Regulation of Mitochondrial Functions in Mice with Shank3 Mutation Associated with Autism Spectrum Disorder. Brain Sci, 11(6). 10.3390/brainsci11060677

Kashiwaya, Y., Takeshima, T., Mori, N., Nakashima, K., Clarke, K., & Veech, R. L. (2000). D-beta-hydroxybutyrate protects neurons in models of Alzheimer’s and Parkinson’s disease. Proc Natl Acad Sci U S A, 97(10), 5440–5444. 10.1073/pnas.97.10.5440

Klimova, N., Long, A., & Kristian, T. (2019). Nicotinamide mononucleotide alters mitochondrial dynamics by SIRT3-dependent mechanism in male mice. J Neurosci Res, 97(8), 975–990. 10.1002/jnr.24397

Lee, Kapur, M., Li, M., Choi, M. C., Choi, S., Kim, H. J.,…Yao, T. P. (2014). MFN1 deacetylation activates adaptive mitochondrial fusion and protects metabolically challenged mitochondria. J Cell Sci, 127(Pt 22), 4954–4963. 10.1242/jcs.157321

Lee, R. W. Y., Corley, M. J., Pang, A., Arakaki, G., Abbott, L., Nishimoto, M.,…Wong, M. (2018). A modified ketogenic gluten-free diet with MCT improves behavior in children with autism spectrum disorder. Physiol Behav, 188, 205–211. 10.1016/j.physbeh.2018.02.006

Lee, S., Chun, H. S., Lee, J., Park, H. J., Kim, K. T., Kim, C. H.,…Kim, W. K. (2018). Plausibility of the zebrafish embryos/larvae as an alternative animal model for autism: A comparison study of transcriptome changes. PLoS One, 13(9), e0203543. 10.1371/journal.pone.0203543

Lee, Y., Ryu, J. R., Kang, H., Kim, Y., Kim, S., Zhang, Y.,…Han, K. (2017). Characterization of the zinc-induced Shank3 interactome of mouse synaptosome. Biochem Biophys Res Commun, 494(3-4), 581–586. 10.1016/j.bbrc.2017.10.143

Lin, A., Turner, Z., Doerrer, S. C., Stanfield, A., & Kossoff, E. H. (2017). Complications During Ketogenic Diet Initiation: Prevalence, Treatment, and Influence on Seizure Outcomes. Pediatr Neurol, 68, 35–39. 10.1016/j.pediatrneurol.2017.01.007

Liu, C. X., Li, C. Y., Hu, C. C., Wang, Y., Lin, J., Jiang, Y. H.,…Xu, X. (2018). CRISPR/Cas9-induced shank3b mutant zebrafish display autism-like behaviors. Mol Autism, 9, 23. 10.1186/s13229-018-0204-x

Lopaschuk, G. D., Karwi, Q. G., Ho, K. L., Pherwani, S., & Ketema, E. B. (2020). Ketone metabolism in the failing heart. Biochim Biophys Acta Mol Cell Biol Lipids, 1865(12), 158813. 10.1016/j.bbalip.2020.158813

Mai, A., Massa, S., Lavu, S., Pezzi, R., Simeoni, S., Ragno, R.,…Sinclair, D. A. (2005). Design, synthesis, and biological evaluation of sirtinol analogues as class III histone/protein deacetylase (Sirtuin) inhibitors. J Med Chem, 48(24), 7789–7795. 10.1021/jm050100l

Miyamoto, J., Ohue-Kitano, R., Mukouyama, H., Nishida, A., Watanabe, K., Igarashi, M.,…Kimura, I. (2019). Ketone body receptor GPR43 regulates lipid metabolism under ketogenic conditions. Proc Natl Acad Sci U S A, 116(47), 23813–23821. 10.1073/pnas.1912573116

Monteiro, P., & Feng, G. (2017). SHANK proteins: roles at the synapse and in autism spectrum disorder. Nat Rev Neurosci, 18(3), 147–157. 10.1038/nrn.2016.183

Napoli, E., Duenas, N., & Giulivi, C. (2014). Potential therapeutic use of the ketogenic diet in autism spectrum disorders. Front Pediatr, 2, 69. 10.3389/fped.2014.00069

Neal, E. G., Chaffe, H., Schwartz, R. H., Lawson, M. S., Edwards, N., Fitzsimmons, G.,…Cross, J. H. (2008). The ketogenic diet for the treatment of childhood epilepsy: a randomised controlled trial. Lancet Neurol, 7(6), 500–506. 10.1016/S1474-4422(08)70092-9

Newman, J. C., & Verdin, E. (2014). beta-hydroxybutyrate: much more than a metabolite. Diabetes Res Clin Pract, 106(2), 173–181. 10.1016/j.diabres.2014.08.009

Newman, J. C., & Verdin, E. (2017). beta-Hydroxybutyrate: A Signaling Metabolite. Annu Rev Nutr, 37, 51–76. 10.1146/annurev-nutr-071816-064916

Oanh, N. T. K., Park, Y. Y., & Cho, H. (2017). Mitochondria elongation is mediated through SIRT1-mediated MFN1 stabilization. Cell Signal, 38, 67–75. 10.1016/j.cellsig.2017.06.019

Pecorelli, A., Ferrara, F., Messano, N., Cordone, V., Schiavone, M. L., Cervellati, F.,…Valacchi, G. (2020). Alterations of mitochondrial bioenergetics, dynamics, and morphology support the theory of oxidative damage involvement in autism spectrum disorder. FASEB J, 34(5), 6521–6538. 10.1096/fj.201902677R

Rossignol, D. A., & Frye, R. E. (2012). Mitochondrial dysfunction in autism spectrum disorders: a systematic review and meta-analysis. Mol Psychiatry, 17(3), 290–314. 10.1038/mp.2010.136

Ruskin, D. N., Murphy, M. I., Slade, S. L., & Masino, S. A. (2017). Ketogenic diet improves behaviors in a maternal immune activation model of autism spectrum disorder. PLoS One, 12(2), e0171643. 10.1371/journal.pone.0171643

Ruskin, D. N., Svedova, J., Cote, J. L., Sandau, U., Rho, J. M., Kawamura, M., Jr.,…Masino, S. A. (2013). Ketogenic diet improves core symptoms of autism in BTBR mice. PLoS One, 8(6), e65021. 10.1371/journal.pone.0065021

Sabouny, R., & Shutt, T. E. (2020). Reciprocal Regulation of Mitochondrial Fission and Fusion. Trends Biochem Sci, 45(7), 564–577. 10.1016/j.tibs.2020.03.009

Samant, S. A., Zhang, H. J., Hong, Z., Pillai, V. B., Sundaresan, N. R., Wolfgeher, D.,…Gupta, M. P. (2014). SIRT3 deacetylates and activates OPA1 to regulate mitochondrial dynamics during stress. Mol Cell Biol, 34(5), 807–819. 10.1128/MCB.01483-13

Santra, S., Gilkerson, R. W., Davidson, M., & Schon, E. A. (2004). Ketogenic treatment reduces deleted mitochondrial DNAs in cultured human cells. Ann Neurol, 56(5), 662–669. 10.1002/ana.20240

Scheibye-Knudsen, M., Mitchell, S. J., Fang, E. F., Iyama, T., Ward, T., Wang, J.,…Bohr, V. A. (2014). A high-fat diet and NAD(+) activate Sirt1 to rescue premature aging in cockayne syndrome. Cell Metab, 20(5), 840–855. 10.1016/j.cmet.2014.10.005

Sheng, X., & Cristea, I. M. (2021). The antiviral sirtuin 3 bridges protein acetylation to mitochondrial integrity and metabolism during human cytomegalovirus infection. PLoS Pathog, 17(4), e1009506. 10.1371/journal.ppat.1009506

Shimazu, T., Hirschey, M. D., Newman, J., He, W., Shirakawa, K., Le Moan, N.,…Verdin, E. (2013). Suppression of oxidative stress by beta-hydroxybutyrate, an endogenous histone deacetylase inhibitor. Science, 339(6116), 211–214. 10.1126/science.1227166

Shutt, T. E., & McBride, H. M. (2013). Staying cool in difficult times: mitochondrial dynamics, quality control and the stress response. Biochim Biophys Acta, 1833(2), 417–424. 10.1016/j.bbamcr.2012.05.024

Singh, K., Singh, I. N., Diggins, E., Connors, S. L., Karim, M. A., Lee, D.,…Frye, R. E. (2020). Developmental regression and mitochondrial function in children with autism. Ann Clin Transl Neurol, 7(5), 683–694. 10.1002/acn3.51034

Spilioti, M., Evangeliou, A. E., Tramma, D., Theodoridou, Z., Metaxas, S., Michailidi, E.,…Gibson, K. M. (2013). Evidence for treatable inborn errors of metabolism in a cohort of 187 Greek patients with autism spectrum disorder (ASD). Front Hum Neurosci, 7, 858. 10.3389/fnhum.2013.00858

Stackley, K. D., Beeson, C. C., Rahn, J. J., & Chan, S. S. L. (2011). Bioenergetic Profiling of Zebrafish Embryonic Development. PLOS ONE, 6(9), e25652. 10.1371/journal.pone.0025652

Tang, G., Gutierrez Rios, P., Kuo, S. H., Akman, H. O., Rosoklija, G., Tanji, K.,…Sulzer, D. (2013). Mitochondrial abnormalities in temporal lobe of autistic brain. Neurobiol Dis, 54, 349–361. 10.1016/j.nbd.2013.01.006

Thorsen, M. (2020). Oxidative stress, metabolic and mitochondrial abnormalities associated with autism spectrum disorder. Prog Mol Biol Transl Sci, 173, 331–354. 10.1016/bs.pmbts.2020.04.018

Triolo, M., & Khacho, M. (2024). Protocol to monitor live-cell, real-time, mitochondrial respiration in mouse muscle cells using the Resipher platform. STAR Protocols, 5(4), 103330. 10.1016/j.xpro.2024.103330

Uddin, G. M., Abbas, R., & Shutt, T. E. (2021). The role of protein acetylation in regulating mitochondrial fusion and fission. Biochem Soc Trans, 49(6), 2807–2819. 10.1042/BST20210798

Valente, A. J., Maddalena, L. A., Robb, E. L., Moradi, F., & Stuart, J. A. (2017). A simple ImageJ macro tool for analyzing mitochondrial network morphology in mammalian cell culture. Acta Histochem, 119(3), 315–326. 10.1016/j.acthis.2017.03.001

Wai, T., & Langer, T. (2016). Mitochondrial Dynamics and Metabolic Regulation. Trends Endocrinol Metab, 27(2), 105–117. 10.1016/j.tem.2015.12.001

Weber, D. D., Aminzadeh-Gohari, S., Tulipan, J., Catalano, L., Feichtinger, R. G., & Kofler, B. (2020). Ketogenic diet in the treatment of cancer - Where do we stand? Mol Metab, 33, 102–121. 10.1016/j.molmet.2019.06.026

Weissman, J. R., Kelley, R. I., Bauman, M. L., Cohen, B. H., Murray, K. F., Mitchell, R. L.,…Natowicz, M. R. (2008). Mitochondrial disease in autism spectrum disorder patients: a cohort analysis. PLoS One, 3(11), e3815. 10.1371/journal.pone.0003815

Whitley, B. N., Engelhart, E. A., & Hoppins, S. (2019). Mitochondrial dynamics and their potential as a therapeutic target. Mitochondrion, 49, 269–283. 10.1016/j.mito.2019.06.002

Xu, F. Y., Taylor, W. A., Hurd, J. A., & Hatch, G. M. (2003). Etomoxir mediates differential metabolic channeling of fatty acid and glycerol precursors into cardiolipin in H9c2 cells. J Lipid Res, 44(2), 415–423. 10.1194/jlr.M200335-JLR200

Yang, H., Sibilla, C., Liu, R., Yun, J., Hay, B. A., Blackstone, C.,…Guo, M. (2022). Clueless/CLUH regulates mitochondrial fission by promoting recruitment of Drp1 to mitochondria. Nat Commun, 13(1), 1582. 10.1038/s41467-022-29071-4

Zhao, Z., Lange, D. J., Voustianiouk, A., MacGrogan, D., Ho, L., Suh, J.,…Pasinetti, G. M. (2006). A ketogenic diet as a potential novel therapeutic intervention in amyotrophic lateral sclerosis. BMC Neurosci, 7, 29. 10.1186/1471-2202-7-29

Zhu, C. T., & Rand, D. M. (2012). A hydrazine coupled cycling assay validates the decrease in redox ratio under starvation in Drosophila. PLoS One, 7(10), e47584. 10.1371/journal.pone.0047584

