## Supplemental Information for "The Ketogenic Diet Metabolite β-Hydroxybutyrate Promotes Mitochondrial Elongation via Deacetylation in HeLa Cells and Improves Autism-like Behavior in Zebrafish"

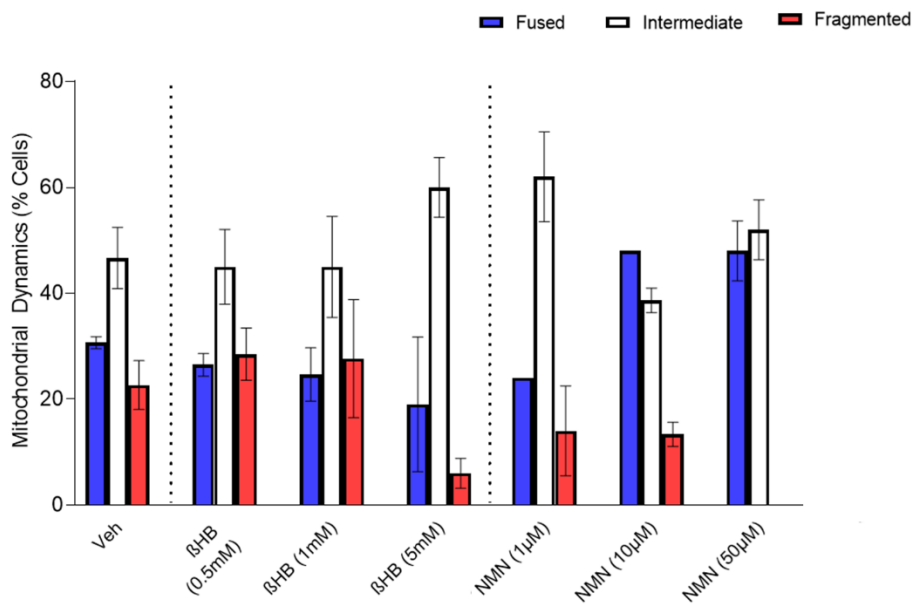

A

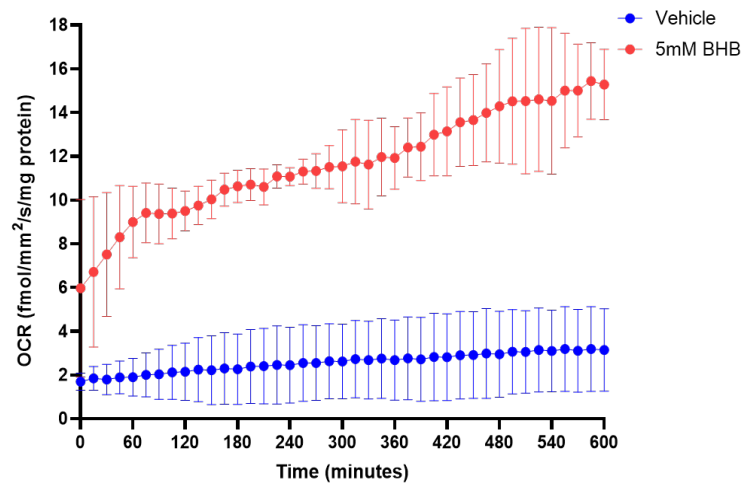

B

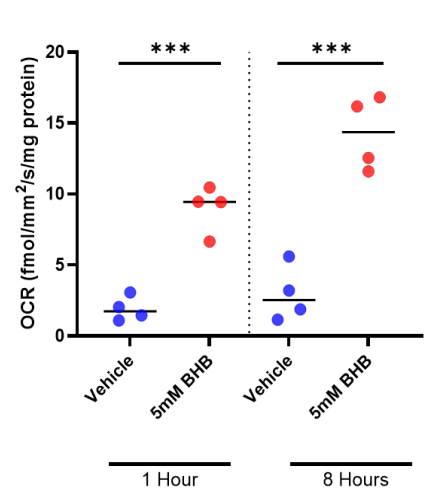

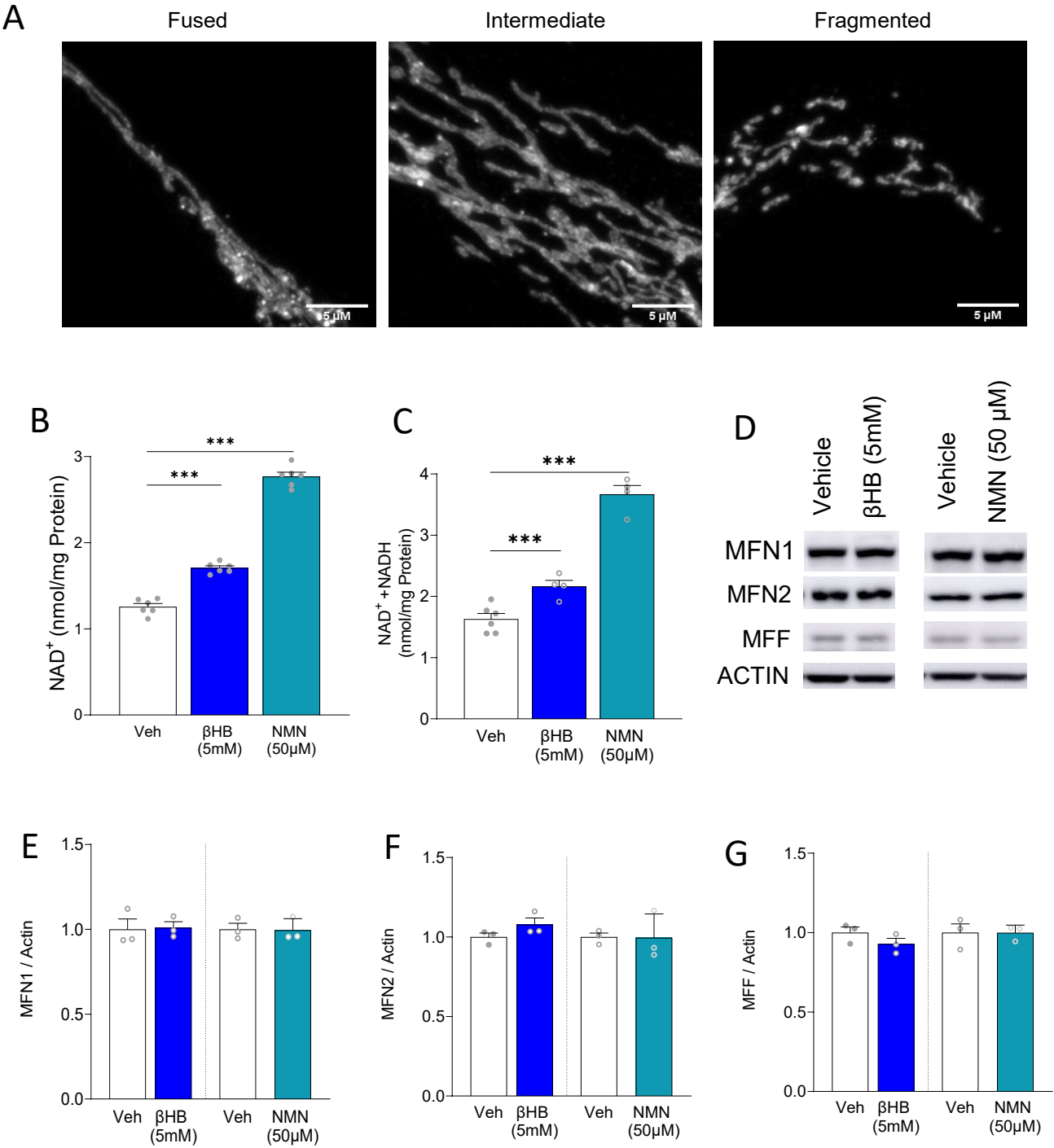

A

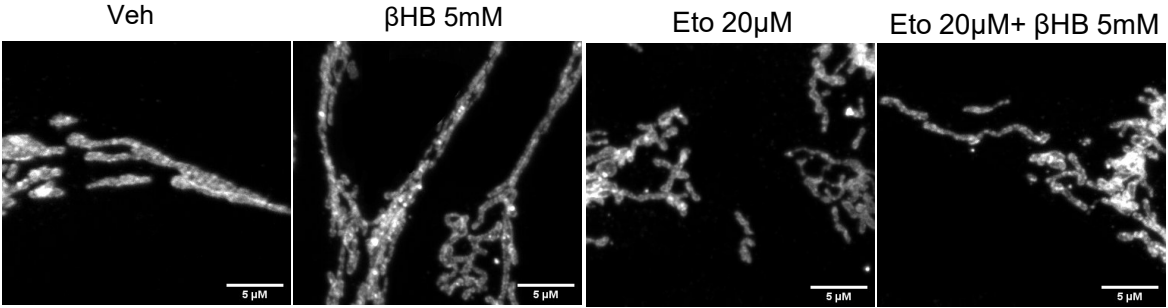

B

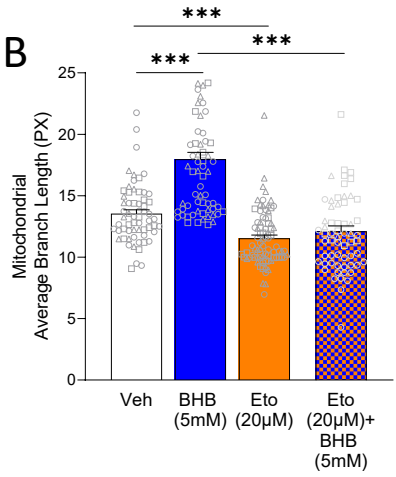

C

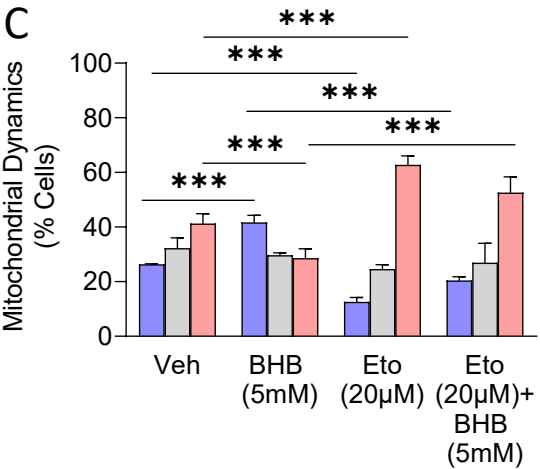

D

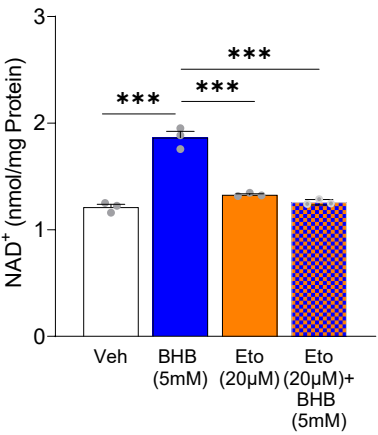

E

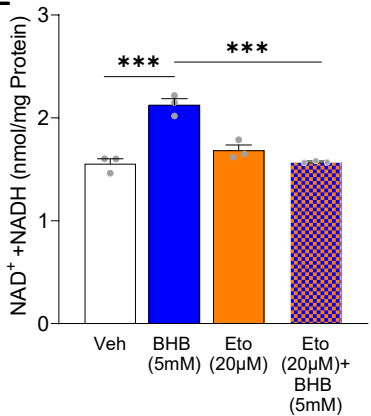

F

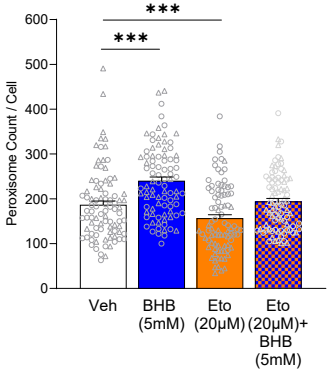

G

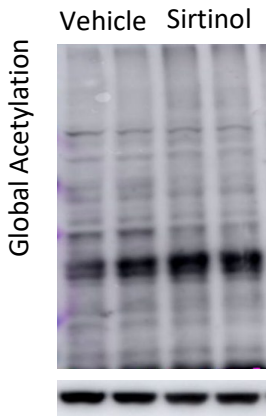

H

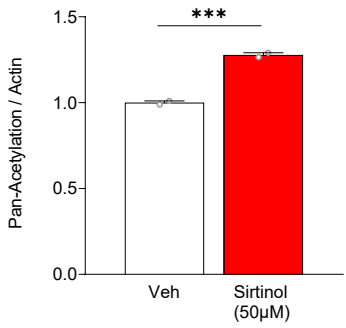

I

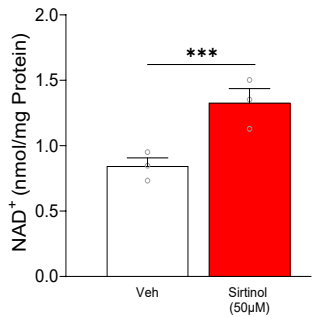

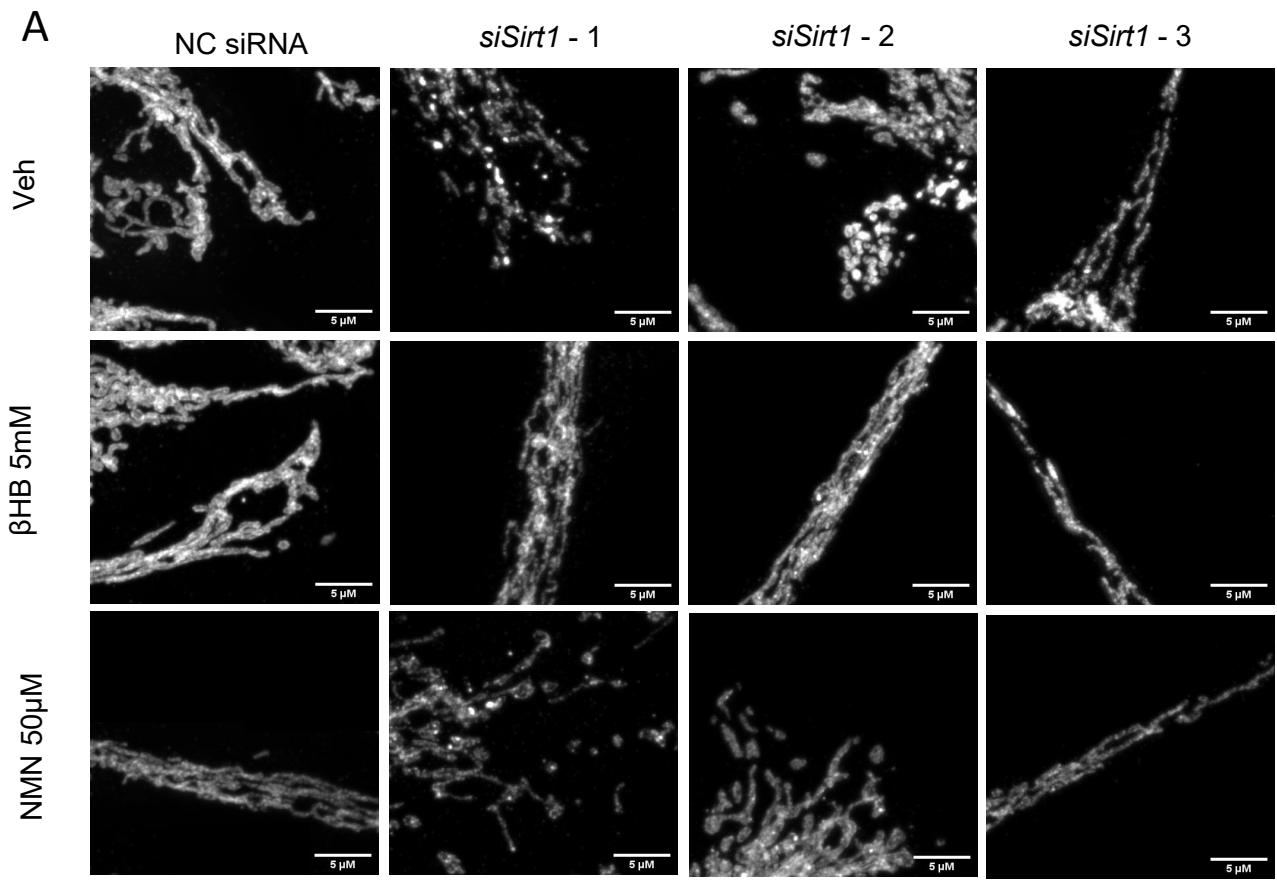

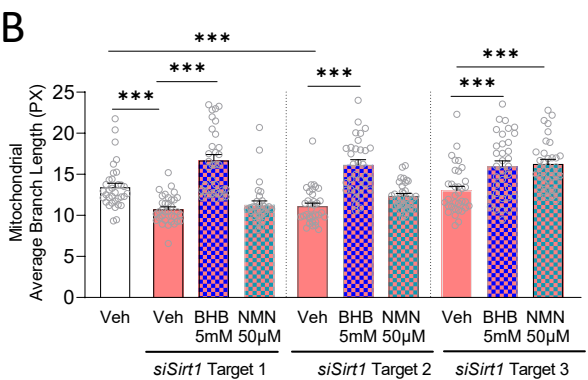

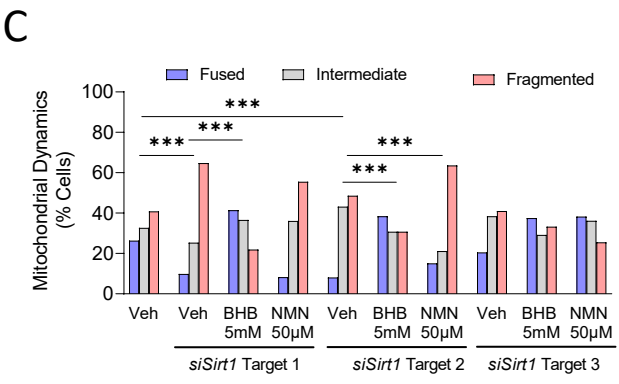

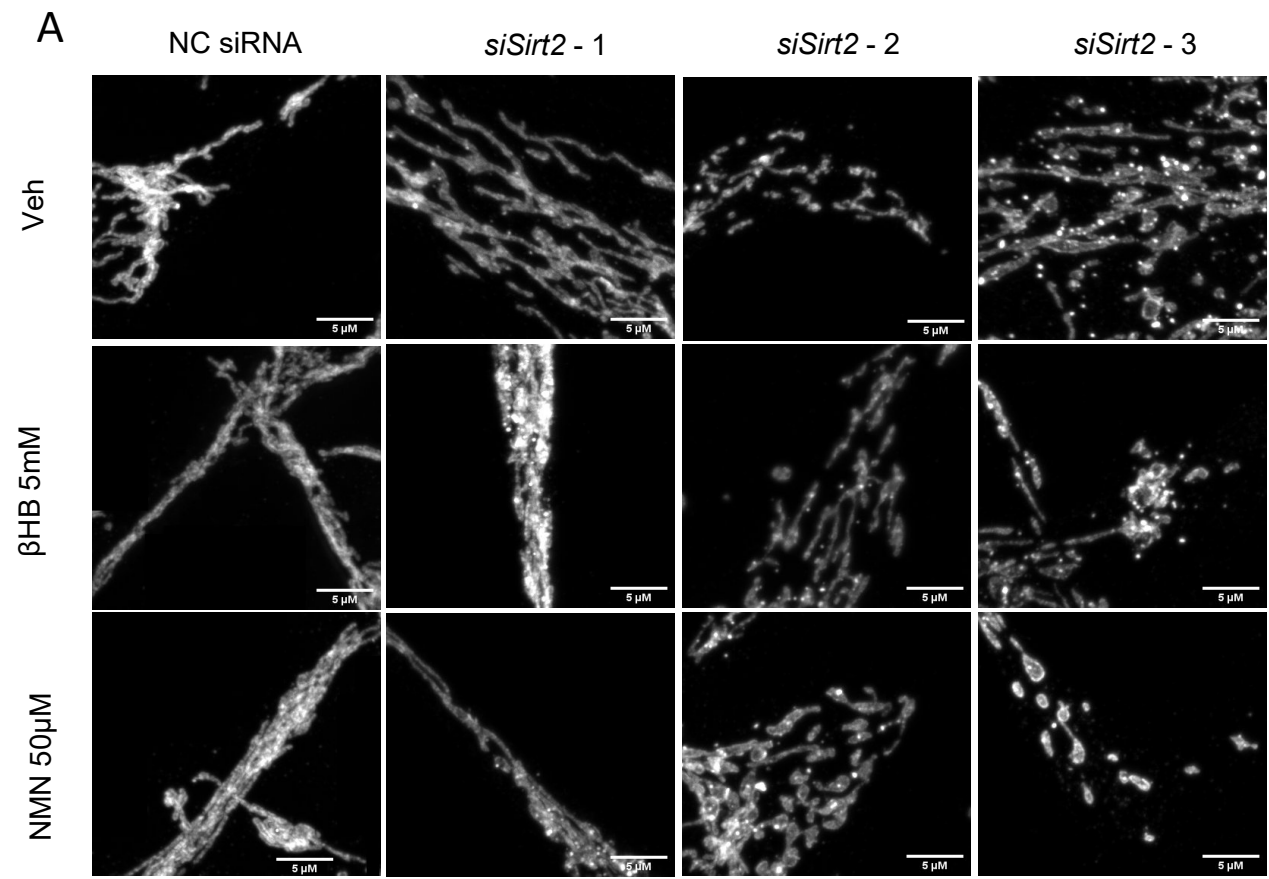

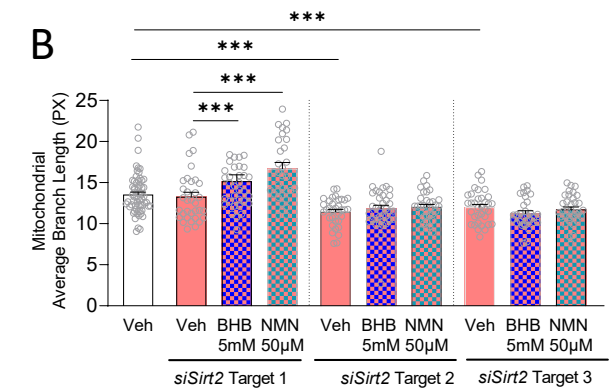

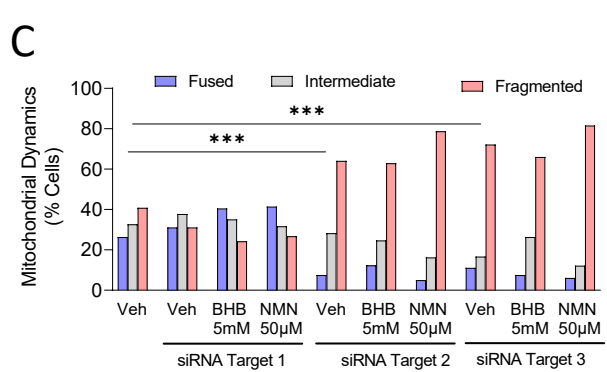

A

NC siRNA

*siSirt3* - 1

*siSirt3* - 2

*siSirt3* - 3

Veh

βHb 5mM

NMN 50μM

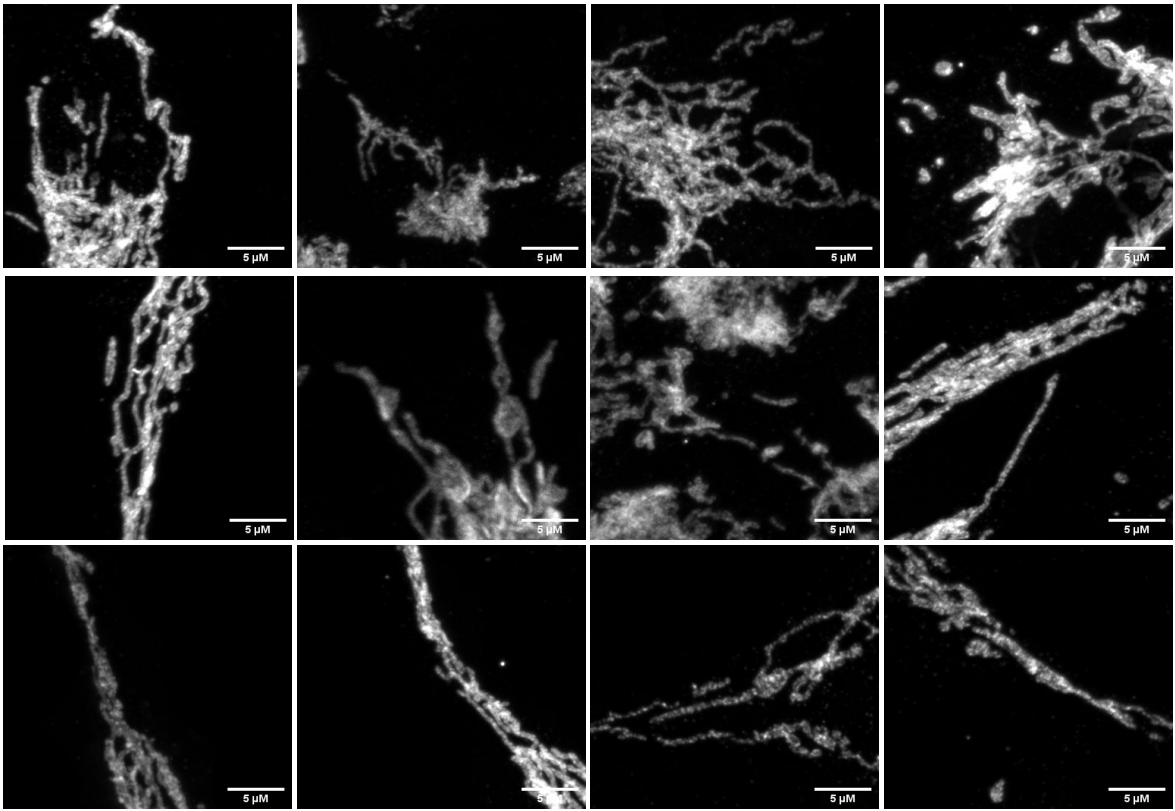

B

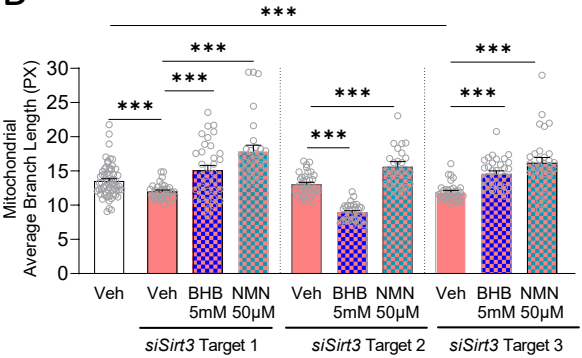

C

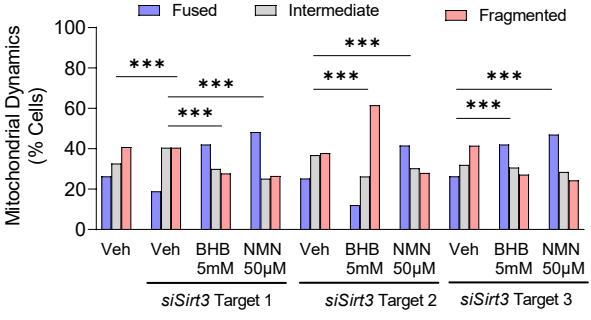

S8A

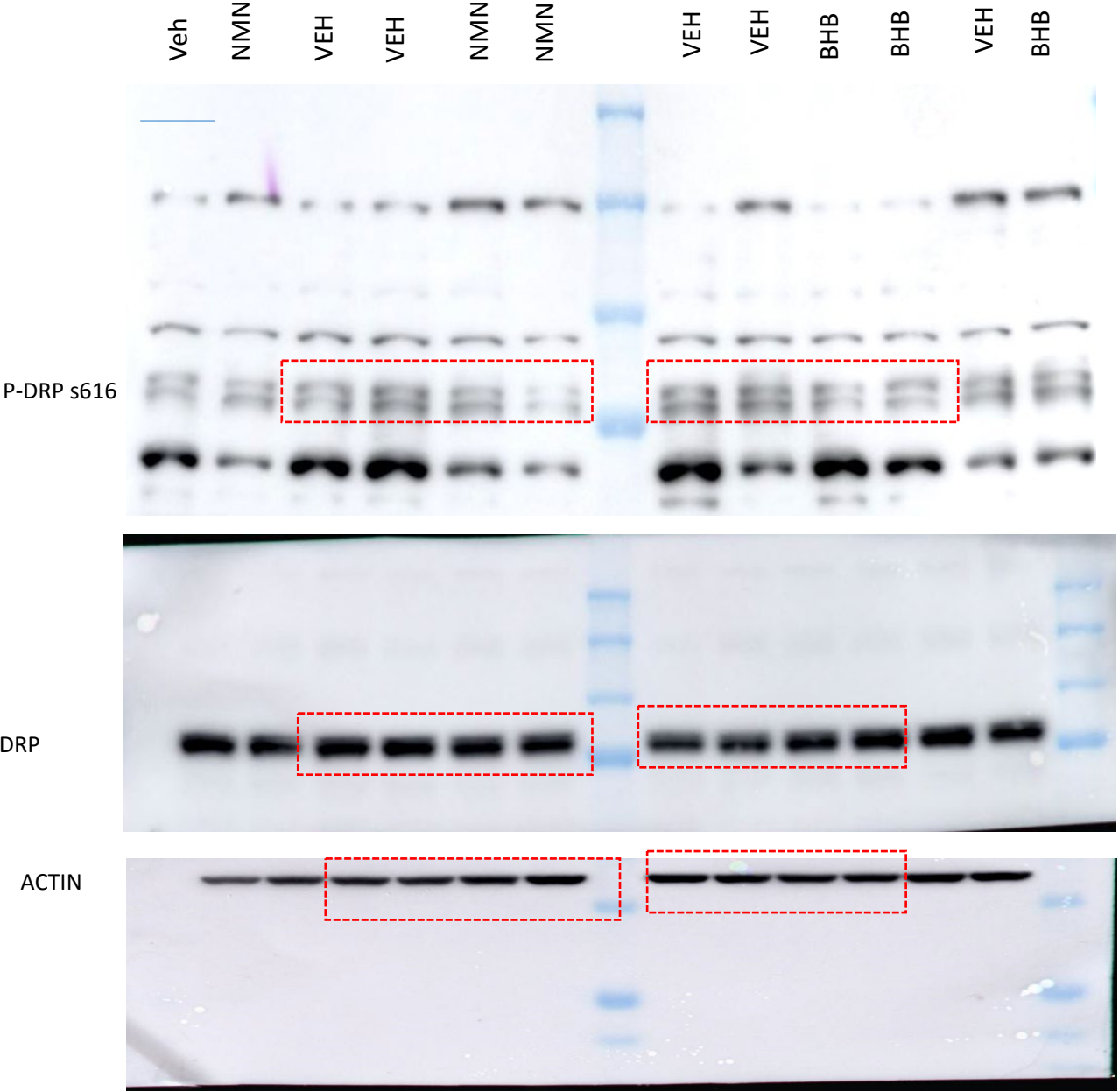

S8B

VEH BHB VEH VEH BHB BHB VEH BHB VEH VEH BHB BHB

OPA1

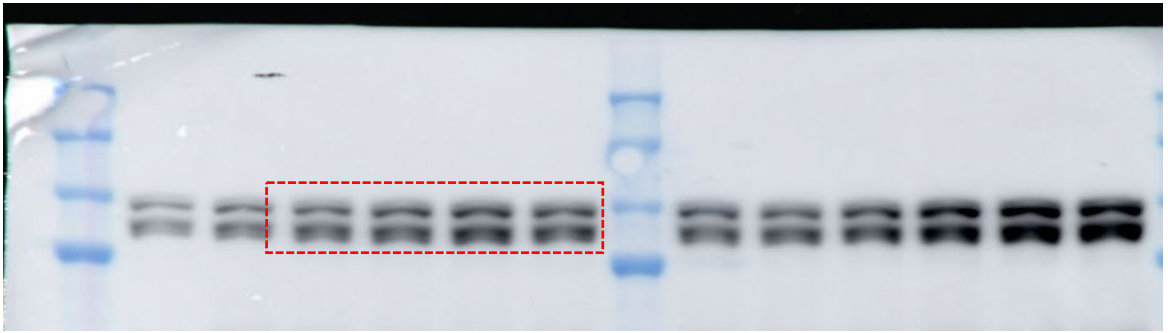

ACTIN

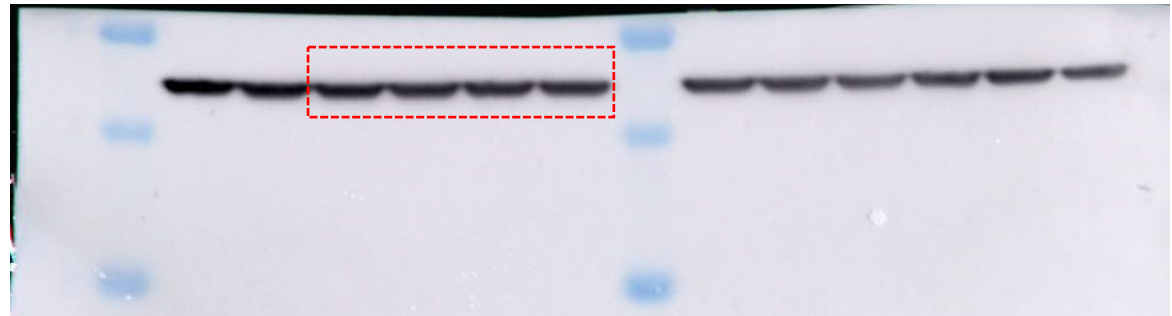

VEH NMN VEH VEH NMN NMN VEH NMN VEH VEH NMN NMN

OPA1

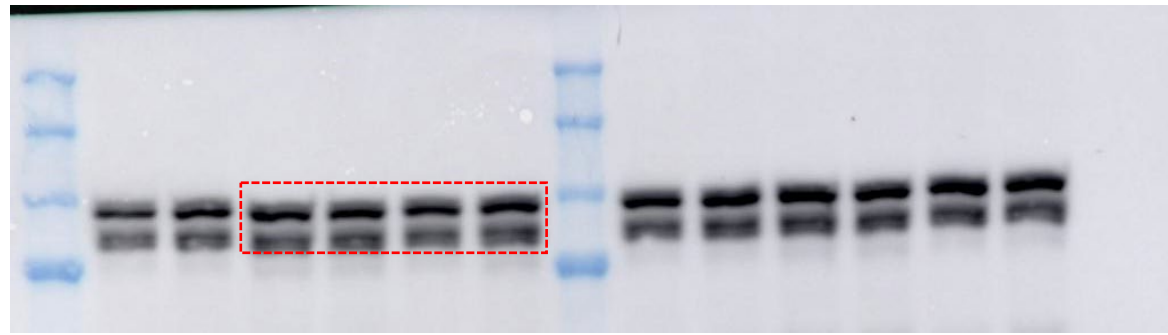

ACTIN

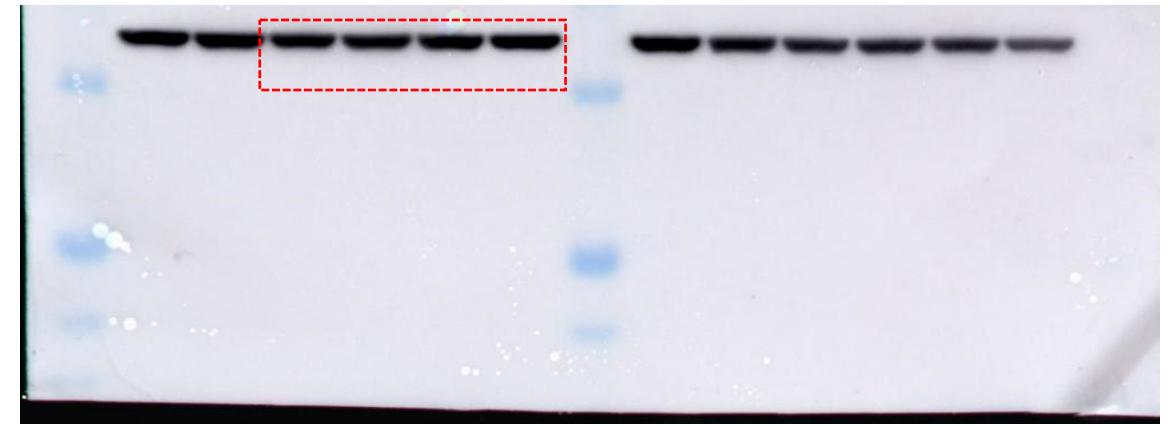

S8C

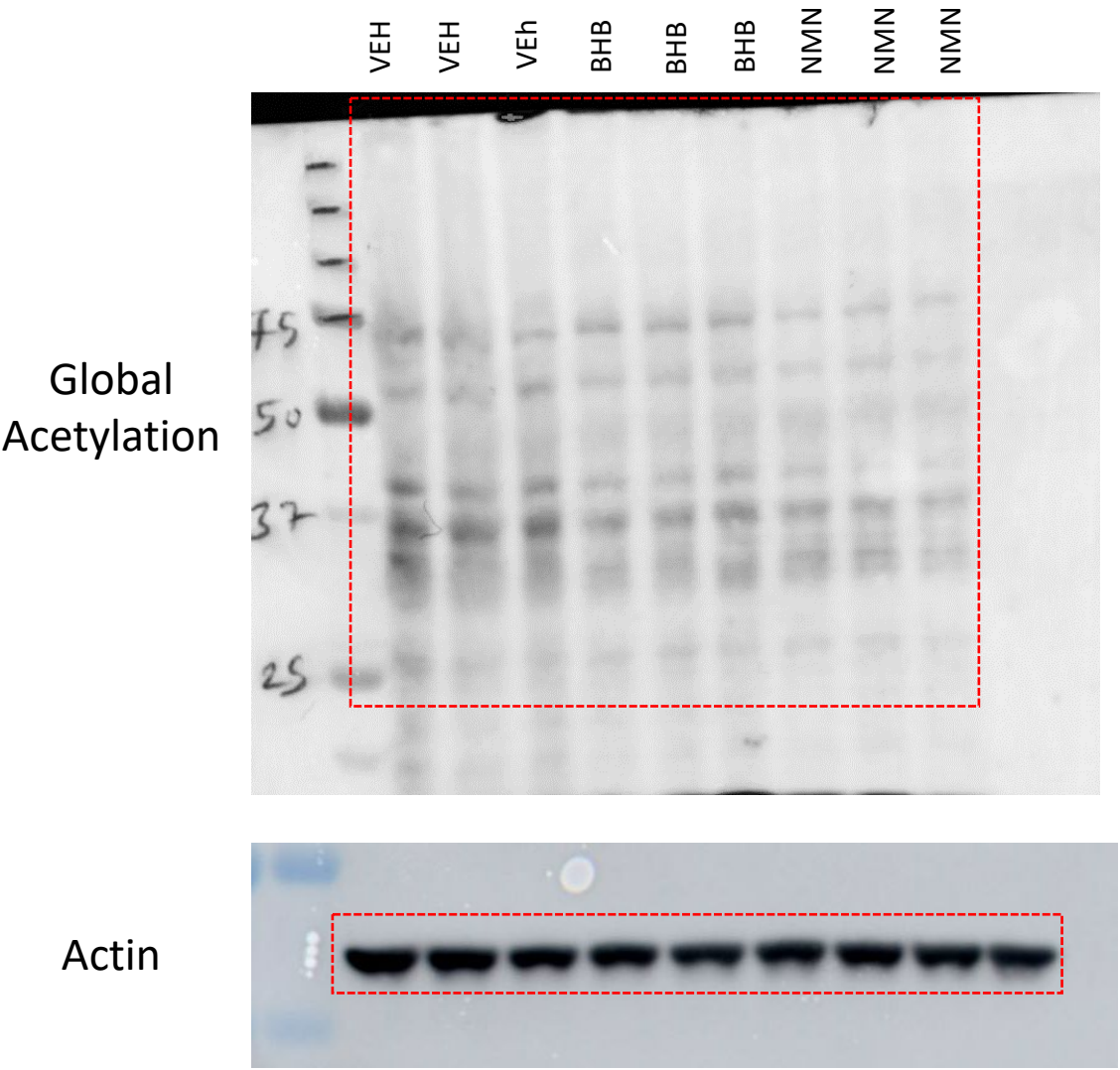

S8D

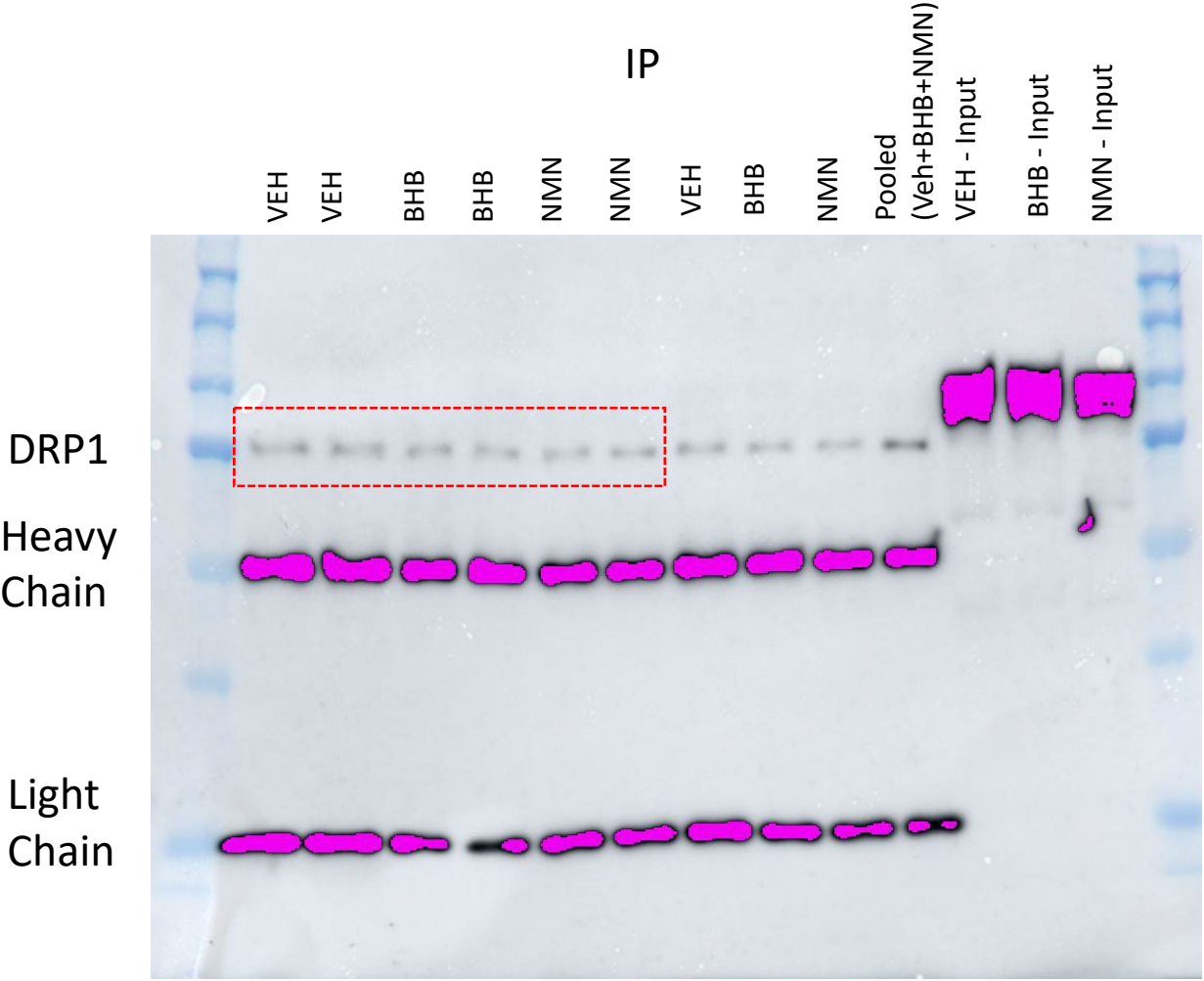

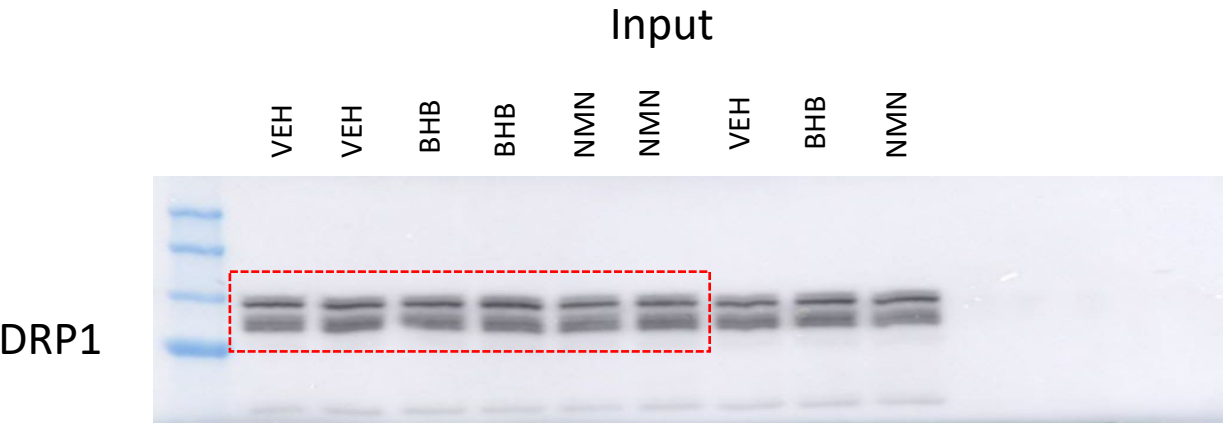

S8E

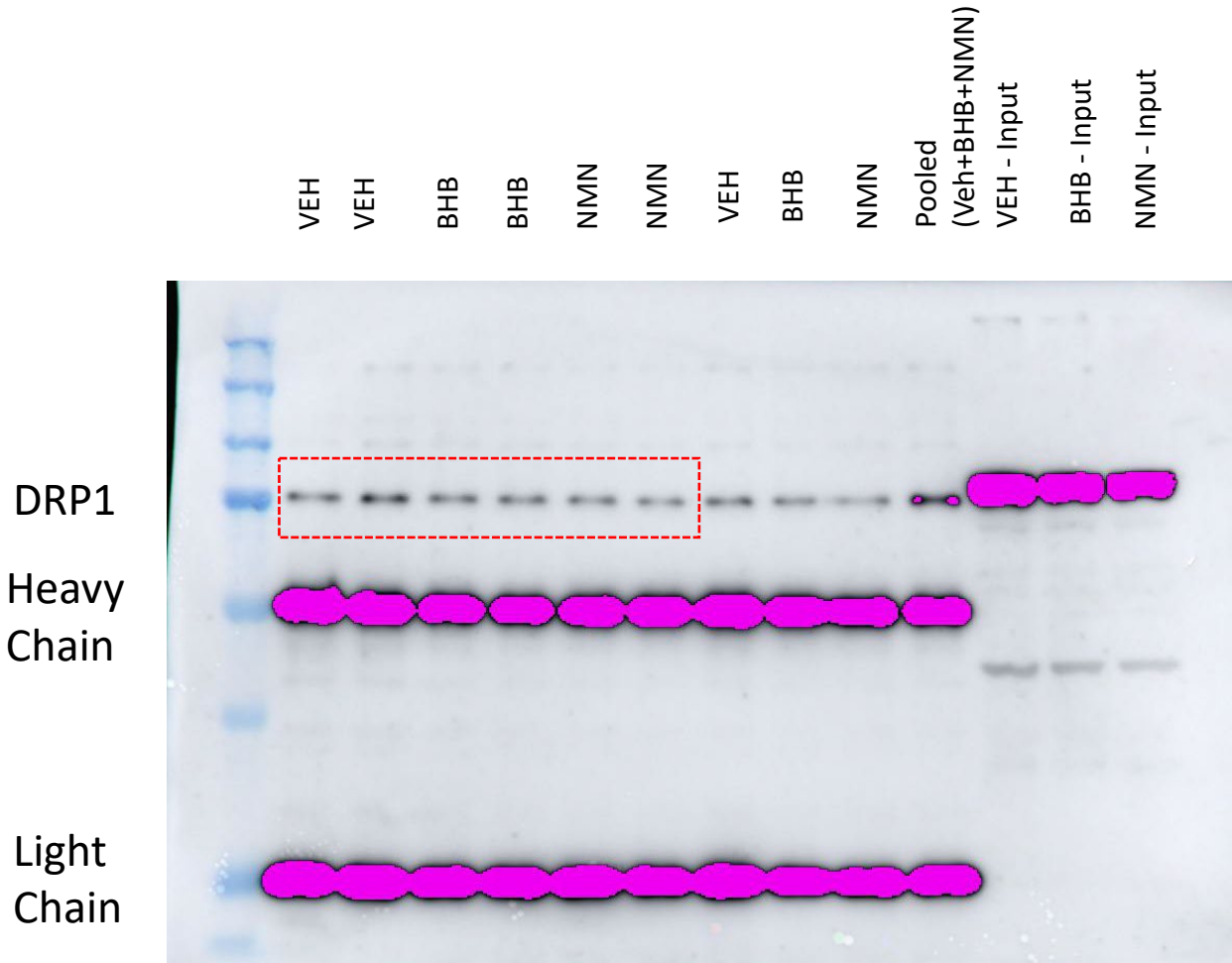

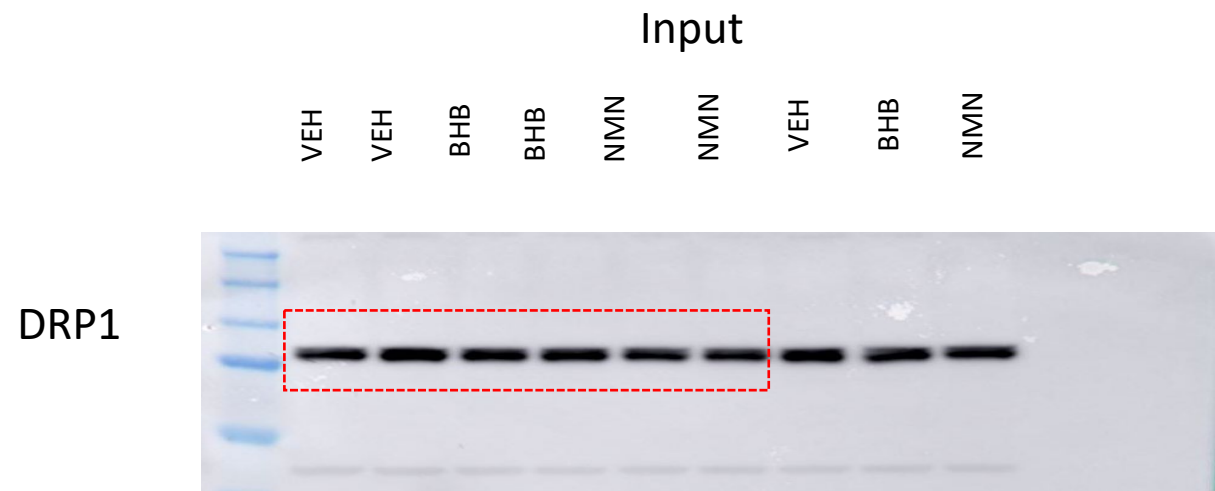

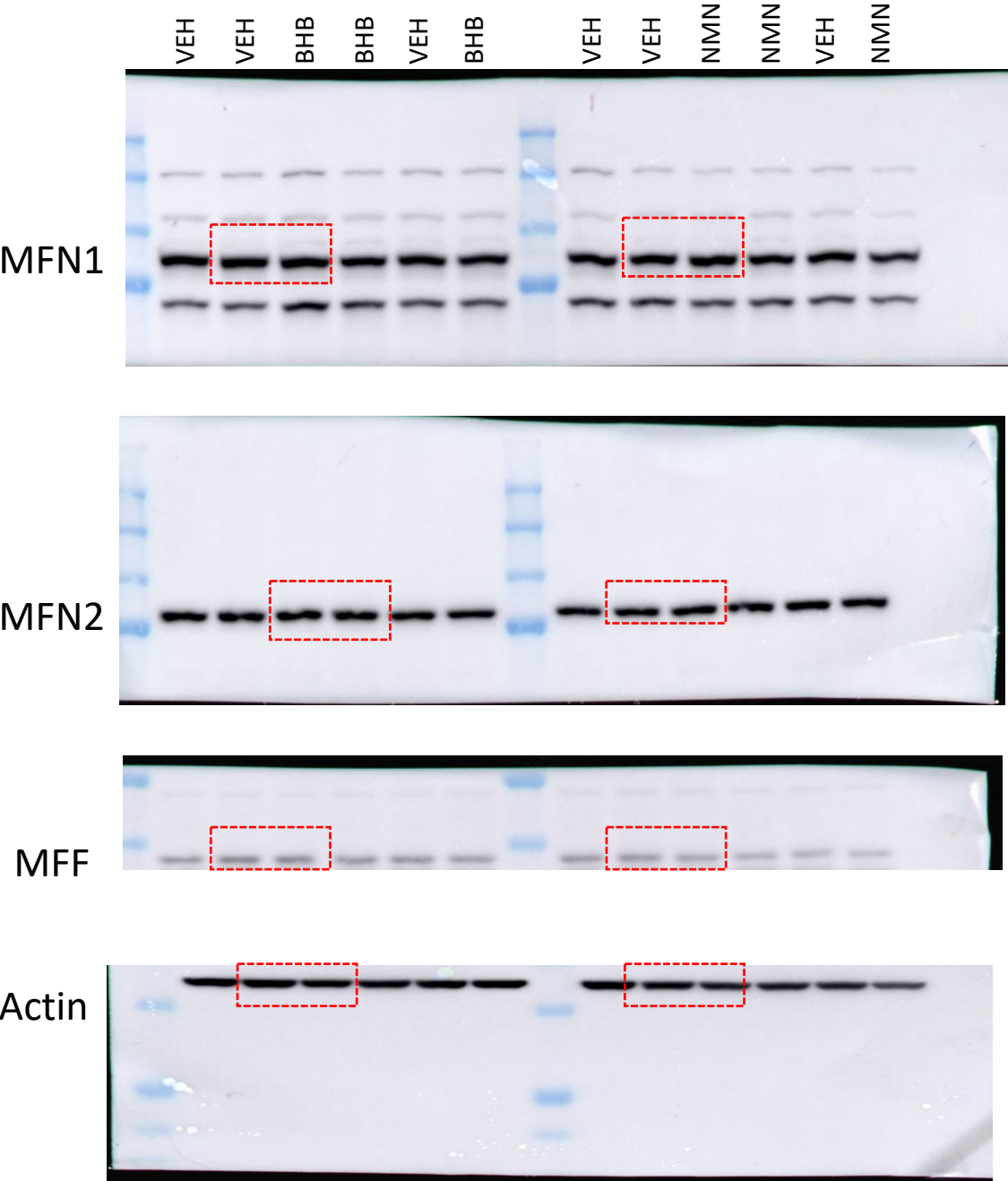
